# RiboTRAP-seq identifies spatially distinct functions for the anterior and posterior intestine in immune and metabolic regulation in *C. elegans*

**DOI:** 10.1101/2025.10.03.680215

**Authors:** Chung-Chih Liu, Nicolas Seban, Supriya Srinivasan

## Abstract

The intestine integrates food-derived cues to coordinate organismal physiology, yet the molecular specialization of discrete regions along the intestinal epithelium remains unclear. Here, we generate cell type-specific translatomes of the *Caenorhabditis elegans* intestine during fasting and refeeding using discrete promoters for the anterior quartet of intestinal cells (INT1) and the remaining 8 pairs of intestinal cells (INT2-9). We found that the anterior-most INT1 cells are a translationally distinct intestinal sub-compartment that is particularly enriched for immune-and stress-response genes. Functional assays using novel INT1-specific genes emerging from this study reveal that these specialized cells play a previously unappreciated role in pathogen avoidance and organismal survival. A second critical function of INT1 cells is their role in sensing and responding to the contents of the gut lumen. We show that luminal pyruvate is the key signal linking bacterial nutrients in the gut, to secretion of the gut insulin antagonist INS-7. These findings establish INT1 as sentinel enteroendocrine cells that integrate metabolic and immune cues to couple food status with immune and endocrine responses. Our studies also provide a rich resource for dissecting segment-specific intestinal biology, an overlooked and fertile area for future research.

## Introduction

The intestine is a critical organ for food digestion and nutrient absorption in animals. *Caenorhabditis elegans*, as a bacterivore, constantly ingests a variety of bacteria with an intestinal transit time as short as two minutes in adults [1]. This rapid process implies the existence of food-sensing mechanisms within the intestine that can continuously monitor and respond to the acute fluctuation in food availability. When food is absent, a transcriptional metabolic response is evoked via the induction of the fasting response genes including lipases, peroxisomal and mitochondrial β-oxidation genes and some fatty acid desaturases in the intestine to maintain energy homeostasis [2–4]. However, it has been unclear whether discrete segments within the intestine have specific food-sensing capabilities or exhibit different responses to fasting in comparison to the organ as a whole. To our knowledge, no studies to date have addressed this question with high spatial and temporal resolution. The *C. elegans* intestine is comprised of 20 cells designated INT1-INT9 [5]. The anterior ring comprises 4 cells called INT1, which are the most proximal intestinal cells adjacent to the pharynx and represent the initial location of contact with ingested food, which for *C. elegans* is digested bacterial components. The remaining 16 cells are arranged in pairs to form the latter 8 posterior rings (INT2-9) arranged in a stereotypical structure. A number of years ago *Sulston et al.* had observed that the microvilli of INT1 cells are shorter than those in the rest of the intestine [5], which implies that INT1 cells may not have the same food absorption capacity that INT2-9 cells have.

In addition to the morphological differences, we recently reported that INT1 functions as an enteroendocrine cell, and secretes the gut peptide INS-7 shortly after food withdrawal. INS-7 secretion levels increase over the duration of fasting, which are reset by the re-introduction of food. Gut INS-7 acts in the nervous system to suppress the fat loss-stimulating peptide FLP-7, functional ortholog of the *C. elegans* tachykinin peptide [6]. This mechanism allows the stimulation of fat loss to be regulated, in part, by incoming food signals from the gut. Furthermore, *Quintin et al.* showed that worms treated with H_2_O_2_ have an induction of the antioxidant enzyme PRDX-2 specifically in INT1 cells, suggesting that INT1 cells may have additional oxidative stress-sensing functions [7]. Although these lines of evidence have suggested that INT1 cells may be a group of specialized intestinal cells that sense the quality of the ingested food and have distinct fasting responses, many questions remain. First, the molecular features of INT1 cells and the molecular distinctions between INT1 and the rest of the intestine (INT2-9) have not been defined. Second, the role of INT1 cells in modulating physiological responses to changes in food availability remains undetermined. Finally, the differential effects of metabolic challenge via fasting and re-feeding in the subsets of intestinal cells are unclear.

In this study, we aim to address these questions using spatial translatomics. In short, the approach is as follows: actively translating mRNAs are bound to clusters of ribosomes, referred to as polysomes [8, 9]. Translating ribosome affinity purification (TRAP) was developed to isolate cell type-specific polysome-mRNA complexes [10, 11], allowing quantitative analysis of translational regulation [12]. Here, we performed TRAP-Seq using a modified protocol based on *McLachlan et al.* [13], in which RPL-22 was tagged with HA to enable affinity purification of polysome-mRNA complexes, which is followed by RNA sequencing. This approach provided a comprehensive spatially distinct translatomic profile of actively translating mRNAs in INT1 and INT2-9 cells. To discern novel temporal features in response to metabolic challenge, we examined the translational responses of these intestinal subsets to acute and chronic fasting and refeeding paradigms. Our datasets reveal that INT1 cells exhibit a distinct molecular identity, characterized by the enrichment of immune- and stress-responsive genes. Furthermore, fasting and refeeding elicit divergent regulatory programs in INT1 and INT2-9 cells. To our knowledge, this represents the first study to determine the molecular identities of discrete intestinal cell subsets in response to fasting and refeeding in *C. elegans*.

## Results

### Isolation and sequencing of translating RNA from INT1 and INT2-9 cells

To identify the molecular differences between INT1 and INT2-9 cells in fed, fasted and refed states, we employed translating ribosome affinity purification (TRAP) [10, 11] to isolate polysomal RNA from INT1 and INT2-9 cells followed by RNA sequencing. This approach captures actively translating mRNAs, thereby providing a cell type-specific snapshot of the translational landscape under fasting and refeeding conditions. To selectively target INT1 and INT2-9 cells, we expressed the HA-tagged ribosomal protein (RPL-22-3xHA) under the control of the *Pges-1ΔB* [6, 14] and *Ppho-1* [15] promoters, respectively. As published [6], we noted that *Pges-1ΔB* is strictly expressed in INT1 cells, whereas *Ppho-1* is expressed in INT2-9 cells with detectable, albeit lower expression in INT2 cells (Fig 1A).

**Fig 1.**
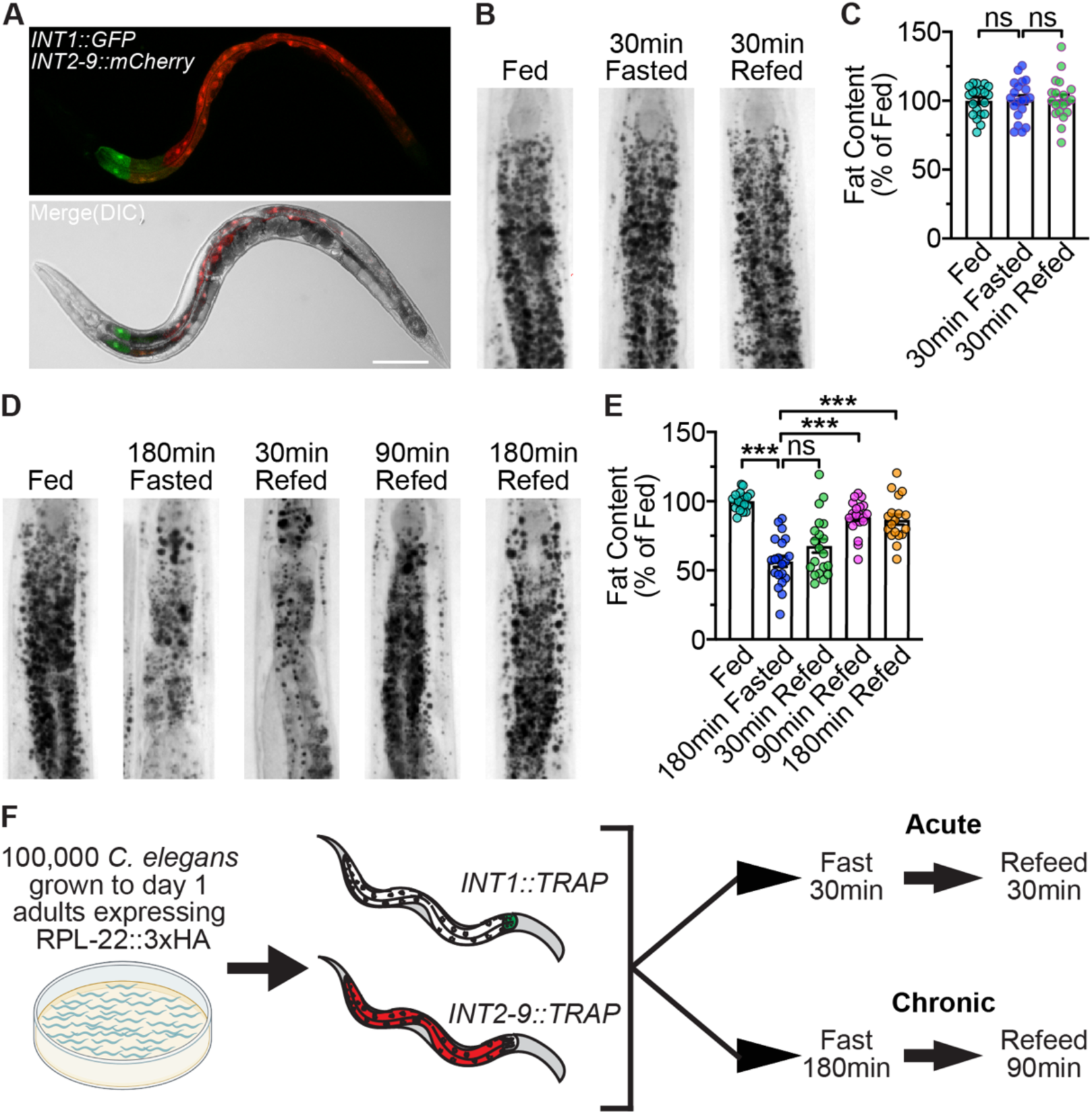
Conditions and workflow for INT1- and INT2-9-specific translatomics. (A) Representative images of wild-type animals bearing integrated *INT1::GFP* and *INT2-9::mCherry* transgenes. Upper panel, GFP expression in INT1 cells and mCherry expression in INT2-9 cells; lower panel, merge with DIC. Scale bar, 100 μm. (B) Representative images of wild-type animals in fed, 30 min fasted, and 30 min refed states fixed and stained with Oil Red O. (C) Fat content was quantified for each condition and expressed as a percentage of wild-type animals in fed state ± SEM (n=20). ^ns^p>0.05 by one-way ANOVA (Tukey). (D) Representative images of wild-type animals in fed, 180 min fasted, 30 min refed, 90 min refed, and 180 min refed states fixed and stained with Oil Red O. (E) Fat content was quantified for each condition and expressed as a percentage of wild-type animals in fed state ± SEM (n=18-20). ***p<0.001, ^ns^p>0.05 by one-way ANOVA (Sidak). (F) 100000 *C. elegans* expressing *INT1::RPL-22::3xHA* or *INT2-9::RPL-22::3xHA* were grown to day 1 adults. Worms were fasted and refed for the time indicated and harvested for TRAP-Seq.

We operationally defined the ‘acute’ conditions at 30 min fasting followed by 30 min refeeding, because under these conditions, intestinal fat content remains unchanged (Fig 1B and 1C). However, INS-7 secretion was induced within the 30 min acute fasting period and restored to baseline levels within 30 min upon refeeding [6], indicating that intestinal responses to food withdrawal can occur before detectable changes in lipid stores. Chronic conditions were established as 3 hr fasting followed by 90 min refeeding. In this paradigm, animals lost approximately half of intestinal lipid stores after chronic fasting, which were subsequently replenished upon refeeding for 90 min (Fig 1D and 1E). Thus, chronic conditions represent a more pronounced metabolic challenge. For TRAP-Seq sample preparation, we cultured 100,000 *INT1::TRAP* or *INT2-9::TRAP* animals to the Day 1 adult stage, subjected them to acute or chronic conditions (Fig 1F), and performed RNA isolation following a modified TRAP protocol [13]. These experiments were conducted 3 times.

### Stress response genes distinguish INT1 from INT2-9 translatomes across acute and chronic conditions

Principal component analysis (PCA) of all TRAP-Seq datasets revealed clearly distinct clustering by promoter identity, indicating that INT1 and INT2-9 cells maintain fundamentally different molecular identities independent of any and all acute or chronic conditions (Fig 2A). We found PC2 to be significantly associated with promoter identity (FDR = 1.2e-10, Pearson’s correlation) and PC1 with metabolic stress (FDR = 1.4e-6, ANOVA). To systematically define these differences, we performed weighted correlation network analysis (WGCNA) [16], which unbiasedly identified 12 gene modules associated with condition-specific responses across the INT1 and INT2-9 datasets (Fig 2B and 2C, S1A Fig). Notably, the expression of genes in module red was positively correlated to all INT1 conditions but negatively correlated to all INT2-9 conditions, suggesting a transcriptional program preferentially translated in INT1 cells (Fig 2C and S1A Fig). Examination of normalized counts confirmed uniformly higher expression of module red genes across all INT1 conditions compared with INT2-9 (Fig 2D). To identify the categories and functions of module red genes, we used WormCat [17, 18], an annotation and category enrichment tool with near-complete annotation of the *C. elegans* genome. Among the 100 genes from module red, we found genes involved in “Stress Response” including C-type lectin, pathogen and heat categories (Fig 2E). In addition, Gene Ontology (GO) term enrichment analysis of module red genes further highlighted the enrichment for the terms “Immune system process” and “Response to biotic stimulus” (S1B Fig).

**Fig 2.**
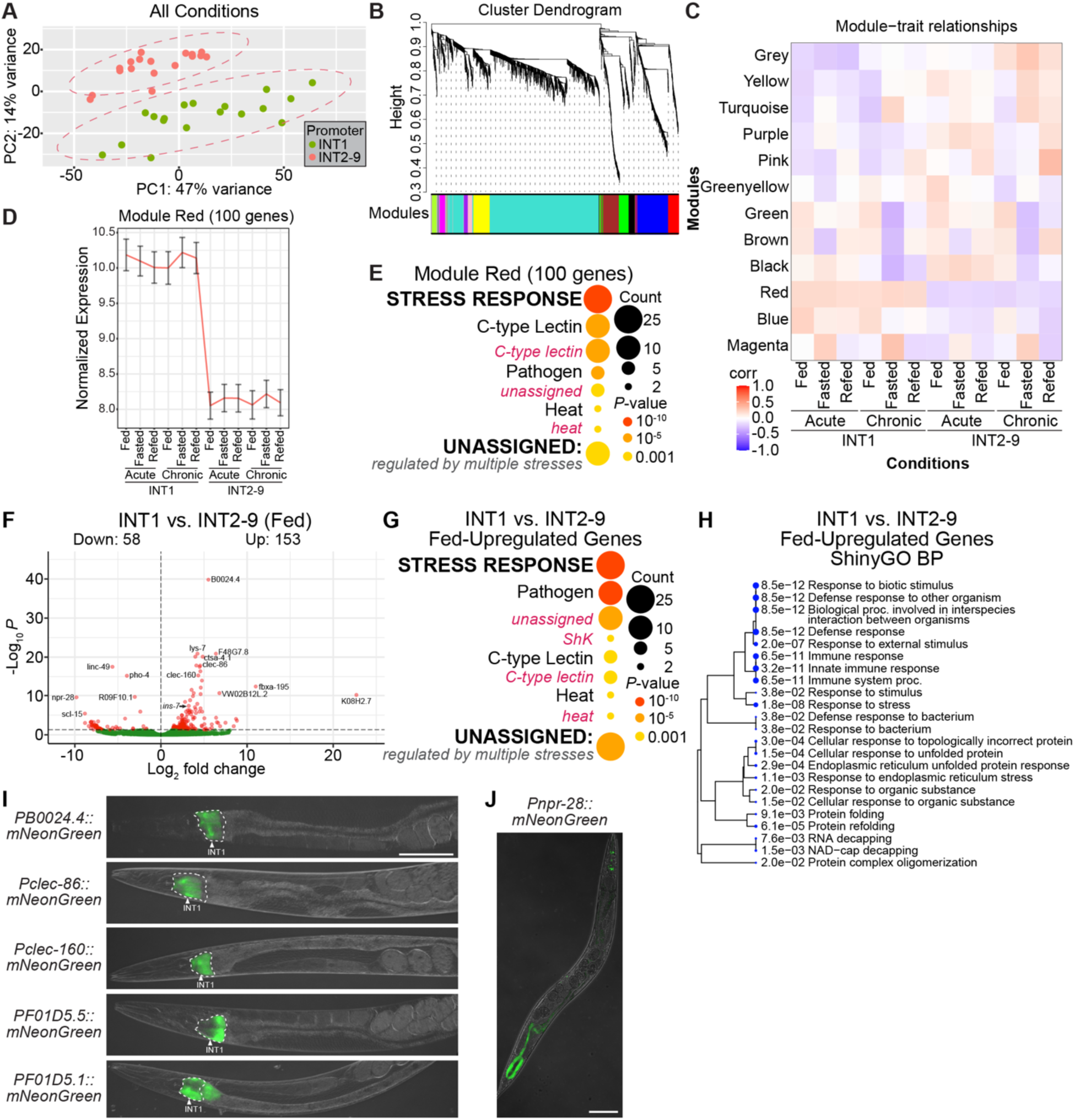
Identification of the fundamental differences between INT1 and INT2-9 translatomes. (A) Principal component analysis showing translatomic variability across the first two principle components of TRAP-Seq data from INT1 and INT2-9 promoters under both acute and chronic conditions (fed, fasted, and refed). Ellipse was computed using the mean and covariance of PC1 and PC2 for each line. (B) Cluster dendrogram produced by average linkage hierarchical clustering in WGCNA. The color row under the dendrogram indicates the module assignment. (C) Pearsons correlations of TRAP-seq conditions with WGCNA modules. Each row represents a WGCNA module and its means correlation to the indicated trait across replicates.The color of each cell indicates the mean pearsons correlation coefficient between the module and the condition; red represents a positive correlation and blue represents a negative correlation. (D) Average expression of red modules genes in the TRAP-seq conditions. The y-axis shows mean variance stabilized expression across all genes and replicates per condition with standard deviation indicated by the error bars. (E) WormCat visualization of categories enriched in genes of module red. Categories 1 are all bold uppercase; Categories 2 are capitalized; Categories 3 are gray italics. The size of the bubbles indicates the gene counts in the category and the color of the bubble represents the adjusted p-value. (F) Volcano plot of differentially expressed genes for INT1 fed versus INT2-9 fed. High confidence differentially expressed genes are labeled. The horizontal dashed line indicates an adjusted p-value cutoff of 0.05. (G) WormCat visualization of categories enriched in genes upregulated in INT1 cells versus INT2-9 cells in the fed state. Categories 1 are all bold uppercase; Categories 2 are capitalized; Categories 3 are gray italics. The size of the bubbles indicates the gene counts in the category and the color of the bubble represents the adjusted p-value. (H) A hierarchical clustering tree of significantly enriched GO terms (Biological Process, BP) in the upregulated genes for INT1 fed versus INT2-9 fed by ShinyGO. GO terms with many shared genes are clustered together. Larger dots indicate more significant adjusted p-values. (I) Fluorescent images of transgenic animals bearing *PB0024.4::mNeonGreen, Pclec-86::mNeonGreen*, *Pclec-160::mNeonGreen*, *PF01D5.5::mNeonGreen or PF01D5.1::mNeonGreen* transgenes. The white arrowhead indicates *mNeonGreen* expression in INT1 cells. Scale bar, 100 μm. (J) Fluorescent image of a transgenic animal bearing *Pnpr-28::mNeonGreen* transgene. Scale bar, 100 μm.

Recently, *Ghaddar et al.* presented a gene expression atlas of adult *C. elegans* with single-cell RNA sequencing [19]. They categorized the intestinal cells into multiple subtypes: intestine anterior (two), intestine middle and intestine posterior (two). To place our findings in the context of previously defined intestinal cell subtypes, we compared the results of our datasets to theirs. INT1 enriched genes were upregulated in the anterior intestine cluster relative to other intestinal subtypes, supporting our conclusion that INT1 represents a translationally distinct intestinal subset (S1C Fig).

Next, we decided to characterize the translatomic differences between INT1 and INT2-9 cells. We began with the differentially expressed genes in INT1 cells relative to INT2-9 cells in the fed state. There were 153 upregulated and 58 downregulated genes (Fig 2F), and we found *ins-7* among the upregulated genes, consistent with our prior report of its restricted expression in INT1 [6]. To identify the categories of the differentially expressed genes in INT1 cells under the fed condition, we performed WormCat category enrichment analysis and also used ShinyGO [20] to conduct Gene Ontology (GO) term enrichment analysis with a hierarchical clustering tree, allowing us to visualize the overlapping relationships among enriched GO terms. In the upregulated genes with WormCat analysis, we found genes involved in “Stress Response” including pathogen, C-type lectin, and heat-shock categories (Fig 2G). Further, the results from ShinyGO demonstrated a cluster of GO terms related to “Immune Response” and “Unfolded Protein Response” (Fig 2H). In contrast, no functional enrichment was detected among the INT1-downregulated genes. Together, these findings show that INT1 cells are characterized by a stress- and immune-responsive translational program in the fed state, marking a strong and clear distinction from the rest of the intestine.

### C-type lectins and ShKT domain-containing proteins are enriched in INT1 cells

We noticed that many INT1-differentially expressed genes were not assigned to a category by WormCat or annotated by ShinyGO. To further explore their molecular functions, we used the InterPro database to annotate the protein domains for each gene and assign the domains into several main categories, such as “Enzyme”, “Membrane Protein”, and “Transcription Factor”. Genes encoding domains that did not fall into these categories, as well as those lacking identifiable domains, were grouped as “Genes with Unknown Functions” (S2A and S2B Fig). Among the 149 upregulated genes in INT1, we found 11 encoding C-type lectins (S2A Fig). C-type-lectin-domain-containing proteins are notable for pathogen recognition and the initiation of immune responses in vertebrates [21]. In *C. elegans,* the expression of many distinct C-type lectins has previously been found to be induced in a pathogen-specific exposure [22, 23].

In addition, we discovered a subset of INT1 upregulated genes encoding ShKT domains within the genes with unknown functions (S2A Fig). The ShK toxin (ShKT) is a potassium (K^+^) channel blocker originally isolated from the sea anemone *Stichodactyla helianthus* [24]. ShKT domain-containing proteins have been reported in parasitic nematode secretomes [25] and shown to modulate the activity of human effector memory T cells by blocking voltage-gated potassium channels [26]. Their biological functions in *C. elegans*, however, remain unclear. Our findings that specific ShKT domain-containing proteins are specifically expressed in INT1 cells are intriguing. Notably, several INT1-upregulated ShKT proteins, including F01D5.1, F01D5.5 and SYSM-1, were previously found to be upregulated by PMK-1 (p38 MAPK) [27], a key regulator of pathogen-induced immune responses in *C. elegans* [27, 28], suggesting that these ShKT proteins may contribute to intestinal immune functions.

To validate our TRAP-Seq findings and define the spatial expression of INT1-enriched genes, we chose key candidate genes with high fold change and low adjusted p-value (therefore, the most significantly altered genes, Fig 2I and S1D-H Fig) to generate transcriptional reporters using their endogenous promoters and 3’UTRs. We started with *B0024.4*, a membrane raft protein-coding gene with the lowest adjusted p-value in the INT1-enriched genes, which revealed tight expression restricted to INT1 cells. We next decided to examine reporters for two C-type lectin genes, *clec-86* and *clec-160*, and two ShKT protein-coding genes, *F01D5.1* and *F01D5.5.* Reporters for *clec-86, clec-160* and *F01D5.5* also exhibited highly specific expression in INT1, whereas *F01D5.1* was expressed in both INT1 and INT2. Out of curiosity, we also generated a reporter line for the most highly downregulated gene *npr-28* in INT1, and interestingly, found that it was restricted to the posterior intestinal cells INT7-9, with no expression in the anterior intestine (Fig 2J). Collectively, these analyses corroborate the spatial restriction of our TRAP-Seq method and demonstrate that INT1-upregulated and INT1-downregulated genes exhibit distinct spatial expression patterns along the intestine.

### Comparisons of enriched categories under acute and chronic conditions identify ShKT domain-containing proteins as stable INT1-specific markers

We next wished to evaluate whether there are specific differences between INT1 and INT2-9 cells under acute fasting and refeeding. In the acute fasted state, INT1 displayed 90 upregulated and 12 downregulated genes (Fig 3A). WormCat analysis of upregulated genes revealed those involved in “Stress Response” including C-type lectin and pathogen categories (Fig 3B), while ShinyGO identified a cluster of GO terms related to “Defense Response” (Fig 3C). In the acute refed state, we observed 52 INT1-upregulated and 35 INT1-downregulated genes (Fig 3D). Enriched categories of the upregulated genes again included “Stress Response”, particularly pathogen and C-type lectins by WormCat (Fig 3E), with GO terms related to “Defense Response” and “Immune Response” by ShinyGO (Fig 3F). InterPro annotations of INT1-upregulated genes revealed consistent representation of “Enzyme”, “C-type Lectin” and “Membrane Protein” categories across fed, acute fasted, and acute refed states (S3A and S3C Fig). Although enriched functional categories were similar between the 2 intestinal sub-compartments, a Venn diagram analysis of the “Stress Response” genes demonstrated only partial overlap across states (Fig. 4A), suggesting that translational differences between INT1 and INT2-9 vary depending on physiological states.

**Fig 3.**
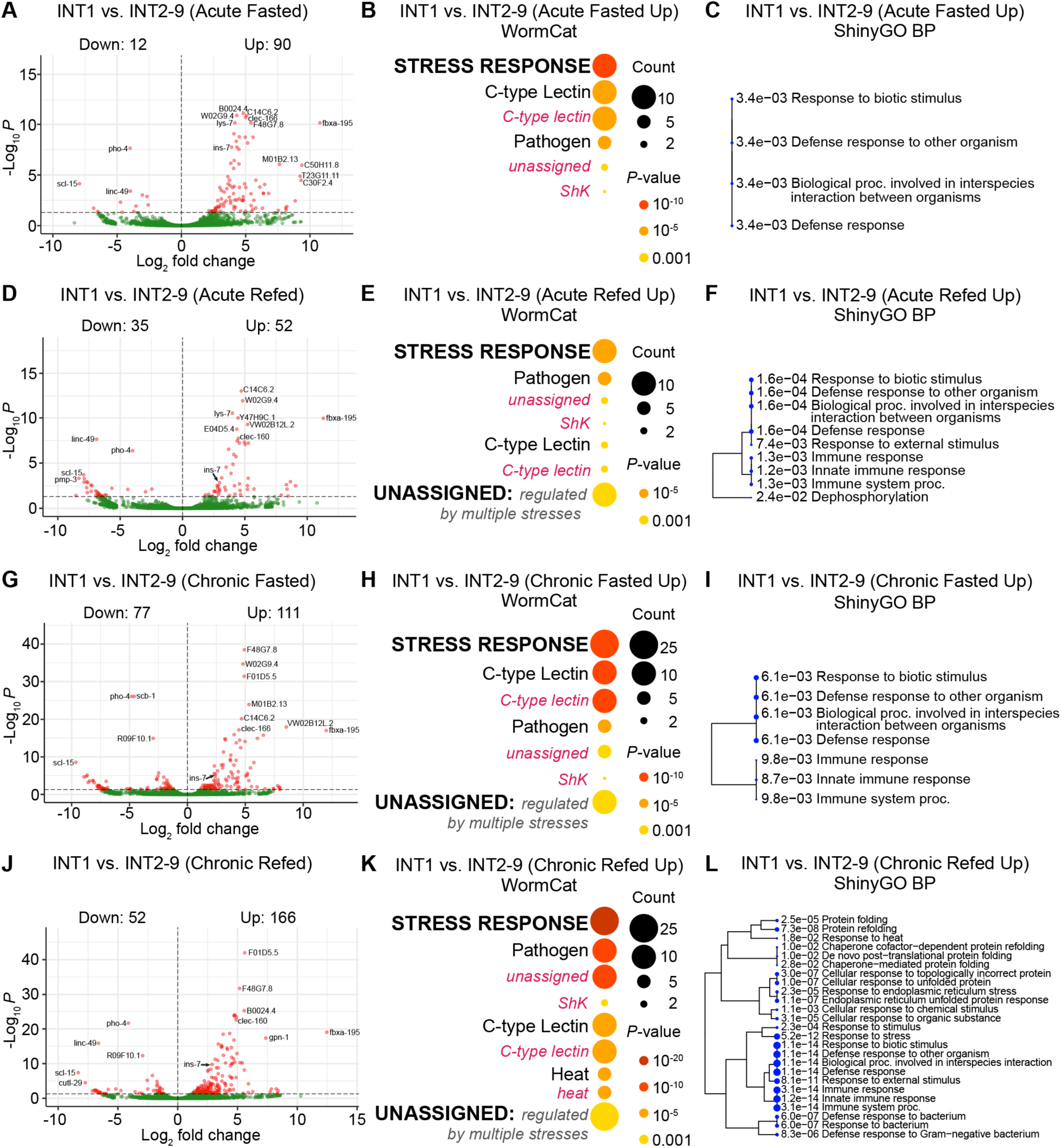
Identification of the differences between INT1 and INT2-9 translatomes under acute and chronic conditions. (A, D, G, J) Volcano plot of differentially expressed genes for INT1 versus INT2-9 in acute fasted, acute refed, chronic fasted and chronic refed states. High confidence differentially expressed genes. The horizontal dashed line indicates an adjusted p-value cutoff of 0.05. (B, E, H, K) WormCat visualization of categories enriched in genes upregulated in INT1 cells versus INT2-9 cells in acute fasted, acute refed, chronic fasted and chronic refed states. Categories 1 are all bold uppercase; Categories 2 are capitalized; Categories 3 are gray italics. The size of the bubbles indicates the gene counts in the category and the color of the bubble represents the adjusted p-value. (C, F, I, L) A hierarchical clustering tree of significantly enriched GO terms (Biological Process, BP) in the upregulated genes for INT1 versus INT2-9 in acute fasted, acute refed, chronic fasted and chronic refed states by ShinyGO. GO terms with many shared genes are clustered together. Larger dots indicate more significant adjusted p-values.

**Fig 4.**
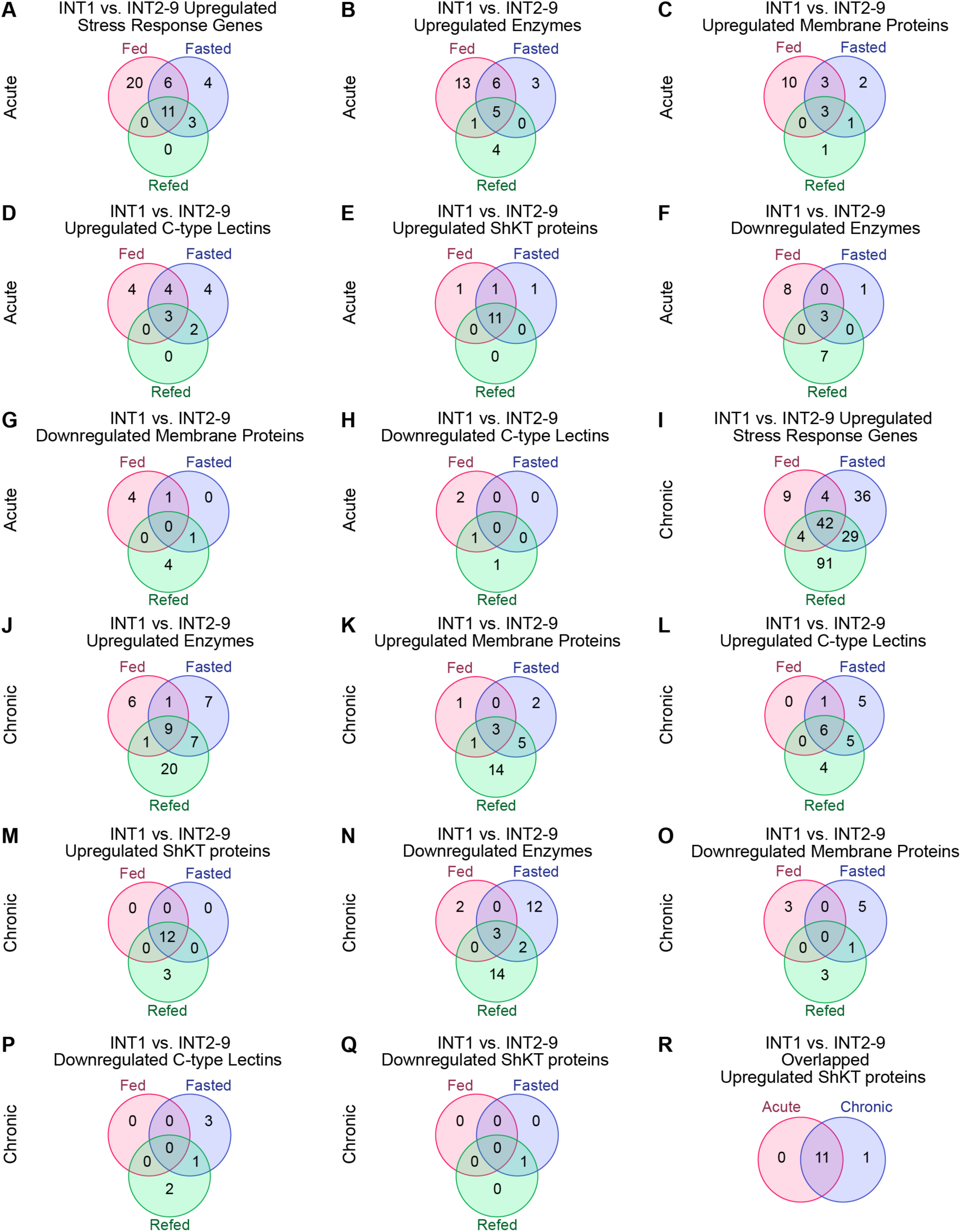
Comparisons of InterPro categories for INT1 versus INT2-9 under acute and chronic conditions. (A-H) Venn diagrams showing the number of upregulated or downregulated genes in specific categories for INT1 versus INT2-9 under the three acute conditions (fed, fasted, and refed). (I-Q) Venn diagrams showing the number of genes in specific categories for INT1 versus INT2-9 under the three chronic conditions (fed, fasted, and refed). (R) Venn diagram showing the overlap between overlapped upregulated ShKT proteins for INT1 versus INT2-9 under acute and chronic conditions.

To further test this possibility, we compared gene lists from 4 major InterPro categories (“Enzyme”, “Membrane Protein”, “C-type Lectin” and “ShKT Protein”) across fed, acute fasted and acute refed states (Fig 4B-H). While “Enzyme”, “Membrane Protein”, and “C-type Lectin” categories exhibited only partial overlap, the “ShKT Protein” category showed near-complete overlap across conditions (Fig 4E), suggesting that ShKT proteins represent stable INT1-specific markers under acute conditions.

We next extended our analysis to chronic fasting and refeeding, which elicit more pronounced metabolic shifts in the intestine. In the chronic fasted state, INT1 exhibited 111 upregulated and 77 downregulated genes (Fig 3G). WormCat analysis of the upregulated genes again identified enrichment for “Stress Response” categories, including C-type lectins and pathogen-related genes (Fig 3H), while ShinyGO revealed a cluster of GO terms related to “Immune Response” (Fig 3I). Similar enrichments were observed in the chronic refed state (Fig 3K and 3L). InterPro annotations of INT1 upregulated genes revealed categories consistent with acute conditions, including “Enzyme”, “Membrane Protein” and “C-type Lectin” (S4A and S4C Fig). Comparison of gene lists demonstrated partial overlap within these categories (Fig 4J-L, 4N-P). Strikingly, however, upregulated ShKT genes showed near-complete overlap across chronic conditions (Fig 4M).

Finally, we compared the INT1-upregulated ShKT genes across both acute and chronic conditions. Venn diagram analysis revealed complete overlap, with the same set of 11 ShKT genes consistently upregulated in all conditions (Fig 4R). These findings demonstrate that ShKT domain-containing proteins represent robust INT1-specific markers, stable across both acute and chronic states. Thus, pathogen-sensing, detoxification and mounting an organismal response to incoming information from the gut lumen are the principal functions of INT1.

### INT1 and INT2-9 cells exhibit distinct fasting and refeeding responses under acute and chronic conditions

Given the fundamentally distinct translatomes of the INT1 and INT2-9 cells, we next investigated whether these differences extend to their responses to fasting and refeeding. Specifically, we sought to determine if the translatomic profiles of INT1 and INT2-9 cells exhibit distinct regulatory patterns in response to metabolic shifts induced by fasting and subsequent refeeding. Looking at the concordance of differentially expressed genes during chronic and acute fasting between INT1 and INT2-9 cells, we identified fasting-response genes with opposing regulation between the two intestinal subsets (Fig 5A and 5B). For instance, 270 genes were upregulated in INT1 following acute fasting but downregulated in INT2-9 under the same condition (Fig 5A), indicating divergent translational responses to metabolic shifts.

**Fig 5.**
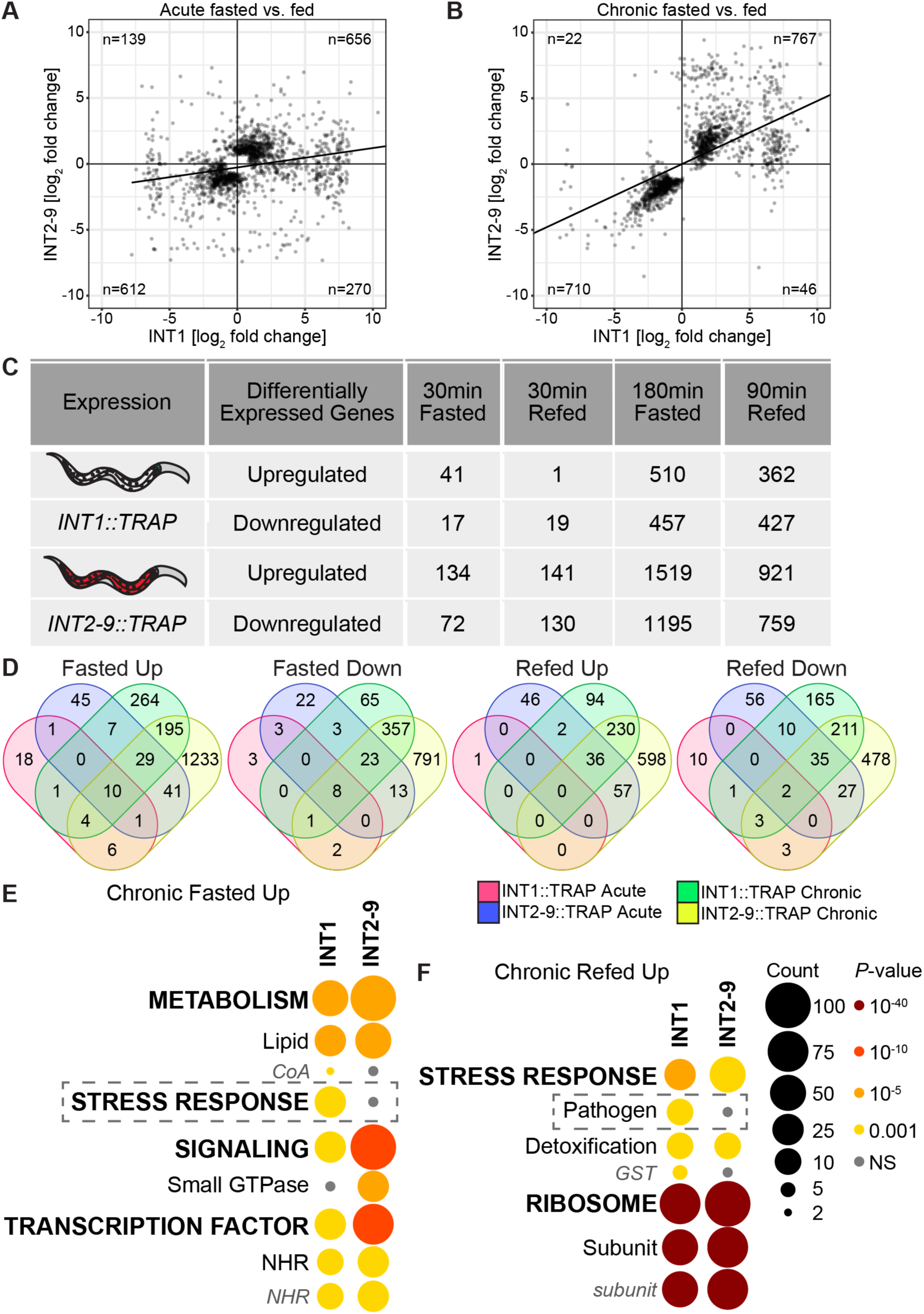
Comparison of fasting and refeeding responses between INT1 and INT2-9 cells under acute and chronic conditions. (A, B) Scatterplots comparing the acute and chronic fasting response genes in INT1 vs INT2-9 differential expression. The top 1000 DEGs by significance were selected for each line and fasting response to be compared, with a regression line fitted to compared INT1 vs INT2-9 response. (C) Table of differentially expressed genes (FDR < 0.05) under acute and chronic conditions for INT1 and INT2-9 datasets. (D) Venn diagrams showing the number of differentially expressed genes (FDR < 0.05) in fasting and refeeding conditions for INT1 and INT2-9 datasets. (E, F) WormCat visualization of categories enriched in upregulated genes in INT1 cells and INT2-9 cells under chronic fasting or refeeding conditions. Categories 1 are all bold uppercase; Categories 2 are capitalized; Categories 3 are gray italics. The size of the bubbles indicates the gene counts in the category and the color of the bubble represents the adjusted p-value. (E) is part of S6A Fig, and (F) is part of S7A Fig.

We next listed all differentially expressed genes (DEGs) in INT1 and INT2-9 under acute and chronic conditions. Overall, INT1 exhibited fewer DEGs than INT2-9 (Fig 5C). To rule out the possibility that INT1 DEGs represented a subset of INT2-9 DEGs, we used Venn diagrams to compare all the INT1 and INT2-9 DEGs from both acute and chronic conditions (Fig 5D). We found two major features of these fasting and refeeding response genes. Notably, INT1 DEGs were not a subset of INT2-9 DEGs regardless of acute or chronic conditions, but rather represented distinct gene sets (Fig 5D). Another key distinction is that for both INT1 and INT2-9, the acute response genes did not overlap substantially with chronic response genes, indicating that distinct translational programs are engaged during acute versus chronic conditions (Fig 5D).

To identify the enriched categories of these INT1 and INT2-9 DEGs under each condition, we performed WormCat analysis and found that INT1 datasets generally had fewer enriched categories than INT2-9 datasets under each condition. (S5A and S5B Fig, S6A and S6B Fig). In the acute fasted state, upregulated genes in both INT1 and INT2-9 were enriched for “Metabolism”, whereas categories including “Stress Response”, “Transcription Factor”, and “Lysosome” were specific to INT2-9 (S5A Fig). Among the downregulated genes in both cell types in the acute fasted state, we found genes involved in “Ribosome”, including ribosomal subunit genes (S5B Fig). Interestingly, in the acute refed state, genes involved in “Ribosome” were upregulated in INT2-9 (S5C Fig), suggesting the sensitivity of ribosomal gene expression to acute fasting and refeeding. Additional categories, such as “Transcription Factor” and “Proteolysis Proteasome”, were enriched exclusively among INT2-9 downregulated genes in the acute refed state (S5D Fig).

Further, we observed that chronic conditions elicited broader and more diverse enrichment patterns than acute states in both INT1 and INT2-9 (S5, S6 and S7 Figs). Although this is an expected result, the category term “Stress Response” was surprisingly enriched only among INT1-upregulated genes in the chronic fasted state (Fig 5E), whereas “Pathogen” was uniquely enriched among INT1-upregulated genes in the chronic refed state (Fig 5F), underscoring an important potential specialization of INT1 in stress and immune responses. Conversely, both INT1 and INT2-9 datasets shared enrichment for “Ribosome” and “Metabolism” among downregulated genes in the chronic fasted state and upregulated genes in the chronic refed state (S6B and S7A Figs), suggesting that protein translation and metabolic regulation are highly responsive to chronic fasting and refeeding.

Taken together, these results demonstrate that INT1 and INT2-9 cells engage distinct translational programs during fasting and refeeding. The divergence is particularly evident under chronic conditions, where INT1 uniquely activates stress- and pathogen-related programs, suggesting specialized roles for INT1 in maintaining homeostasis under chronic conditions. However, because a large portion of the INT1 DEGs remain unassigned by WormCat, the full extent of INT1-specific regulatory programs may not be fully captured by gene set enrichment analyses.

### INT1 cells modulate pathogen avoidance, survival, and peptide secretion

To begin to annotate and assign functions to the genes emerging from our datasets, we examined the pathogen response and peptide secretion functions of the INT1 cells, and the roles of the most highly regulated genes therein. To study pathogen response, we noted that previous studies had implicated INT1-enriched genes in immune responses. For example, expression of *B0024.4* is induced upon exposure to pathogenic *S. maltophilia* JV3 strain [29], and *clec-42,* one of our INT1-upregulated genes, is exclusively expressed in INT1 at all larval stages and mediates resistance to pathogen Bt247 [23]. To test whether INT1-upregulated genes contribute to immune responses, we used an INT1-specific RNAi strain (*sid-1; Pges-1ΔB::sid-1*) to knock down *clec-86* and *clec-160*, and then performed PA14 avoidance and PA14 slow-killing assays [30]. Interestingly, animals with INT1-specific knockdown of either gene exhibited impaired PA14 avoidance (Fig 6A and 6B) and significantly increased survival on PA14 (Fig 6C). These findings suggest that INT1 cells execute specialized, gene-dependent immune functions during pathogen exposure.

**Fig 6.**
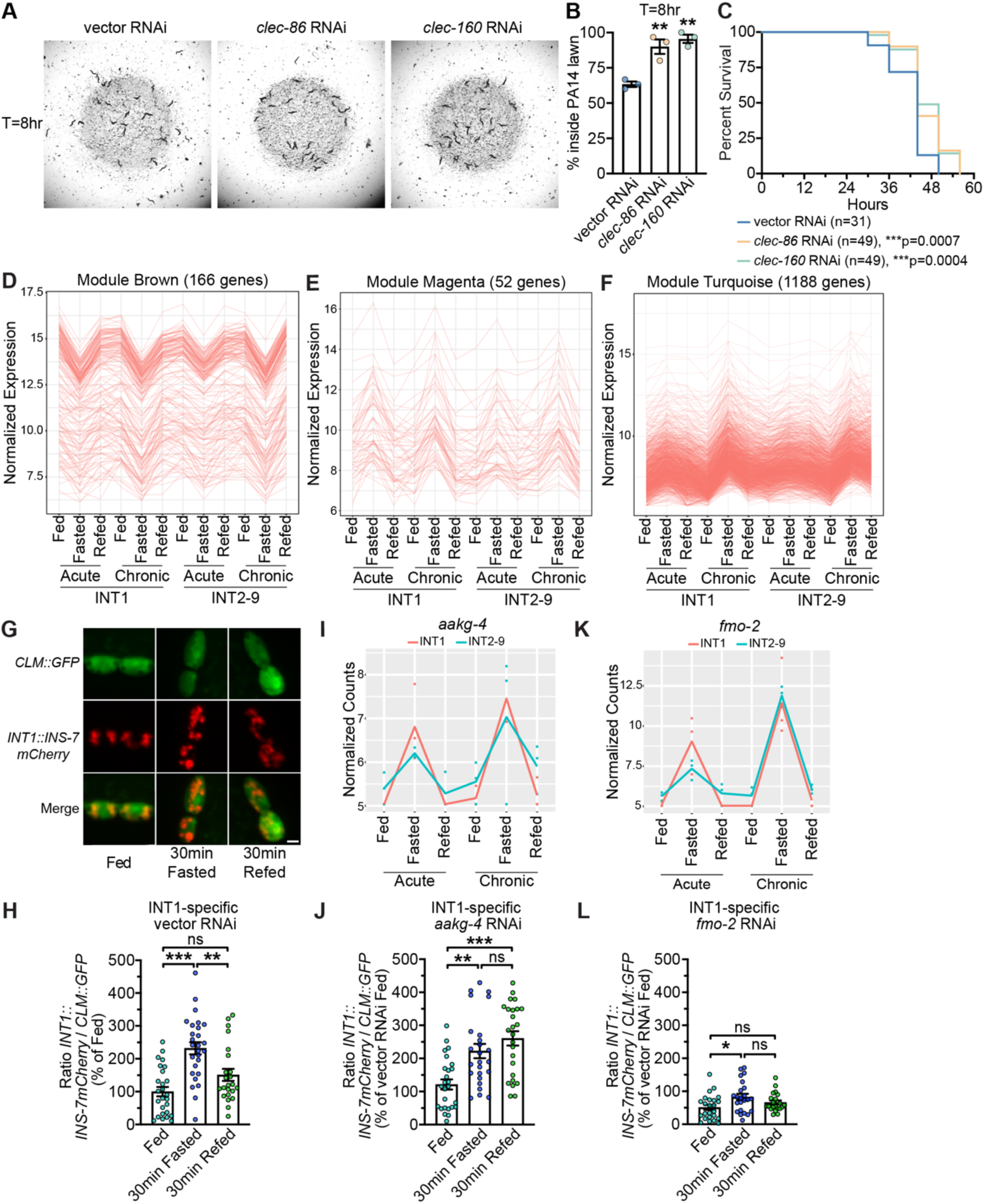
INT1-specific genes modulate behavior and metabolism. (A) The vector, *clec-86*, and *clec-160* RNAi treated *sid-1;INT1::sid-1* animals after 8 hr exposure to PA14 lawns are shown. (B) The distribution of indicated RNAi-treated *sid-1;INT1::sid-1* animals at hour 8 after being transferred to PA14 plates. At hour 0, a total of 30 animals were placed near the PA14 lawn. Data are expressed as a percentage of the animals staying inside PA14 lawn at hour 8 ± SEM (n=3, 3 plates per RNAi treatment, 30 animals on each plate). **p<0.01 by one-way ANOVA (Dunnett). (C) The lifespan of each indicated RNAi-treated animal after being transferred to PA14 lawn was determined by counting the number of live and dead animals every 6hr until all animals had died. Data are plotted as the percentage of animals that survived on any given hour relative to the number of live animals at hour 0. The number of animals for each RNAi treatment is indicated in the figure panels. ***p<0.001 by Log-rank Test. The median survival is 44 hours for all the RNAi treatments. (D-F) Average expression of brown, magenta, and turquoise modules genes in the TRAP-seq conditions. The y-axis shows mean variance stabilized expression across all genes and replicates per condition and module with standard deviation indicated by the error bars. (G) Representative images of *sid-1;INT1::sid-1* animals bearing integrated *INT1::INS-7mCherry* and *CLM::GFP* transgenes treated with vector RNAi at the indicated time points. Scale bar, 5 μm. (H, J, L) *sid-1;INT1::sid-1* animals bearing integrated *INT1::INS-7mCherry* and *CLM::GFP* transgenes were treated with vector, *aakg-4* or *fmo-2* RNAi. INS-7 secretion levels were measured after 30 min fasting and after 30 min refeeding. The intensity of INS-7mCherry fluorescence within a single coelomocyte was quantified and normalized to the area of CLM::GFP expression for each RNAi treatment. Data are expressed as a percentage of the normalized INS-7mCherry fluorescence intensity of fed animals treated with vector RNAi ± SEM (n=21-27). *p<0.05, **p<0.01, ***p<0.001, ^ns^p>0.05 by one-way ANOVA (Dunnett T3). (I, K) Normalized expression plots of *aakg-4* and *fmo-2* under all conditions in INT1 and INT2-9 cells by replicate.

From the WGCNA, we noted that some modules (brown, magenta, turquoise) contained genes with an undulating change in expression levels in response to fasting and refeeding under both acute and chronic conditions (Fig 6D-F). These genes could be classified as “fasted up/refed down” or “fasted down/refed up”. WormCat analysis of brown module genes identified enrichment for “Ribosome” and “Metabolism”, consistent with categories previously linked to ribosomal subunit genes (S6B, S7A, and S8A Figs). Notably, modules magenta and turquoise contained “fasted up/refed down” genes. To determine the strength of the induction across lines we used a Wilcoxan test comparing the distribution of the normalized expression of module genes between fed and fasted conditions. We found the induction in INT2-9 cells (p = 5.6e-4, average LFC = 0.07) after acute fasting was weaker in the turquoise module when compared to INT1 cells (p = 5.6e-50, average LFC = 0.12), suggesting these undulating genes were more INT1-specific (Fig 6E and 6F).

Given our prior finding that the gut peptide INS-7 is secreted from INT1 cells in response to acute fasting [6], we next investigated whether INT1-specific undulating genes could modulate this response. Using the INT1-specific RNAi strain (*sid-1; Pges-1ΔB::sid-1*) crossed into an INS-7 secretion reporter, we confirmed that vector RNAi-treated animals displayed the expected undulating INS-7 secretion levels upon acute fasting and refeeding (Fig 6G and 6H), as previously described [6]. We next targeted two INT1-specific undulating genes, *aakg-4* and *fmo-2*, which showed the largest absolute fold changes across acute fasting and refeeding for INT1-specific RNAi (Fig 6I and 6K). *aakg-4* is the gene encoding one of the five γ subunits of the AMP-activated protein kinase (AMPK) in *C. elegans* [31]. However, the physiological role of *aakg-4* in the intestine is unknown. Remarkably, INT1-specific *aakg-4* knockdown prevented INS-7 secretion from returning to basal levels after refeeding, indicating a requirement for *aakg-4* in resetting INS-7 secretion and suggesting that INS-7 release from INT1 cells is subject to energy-dependent regulation, consistent with its modulation by fasting and refeeding (Fig 6J). The second gene *fmo-2*, encoding flavin-containing monooxygenase-2, is known to promote longevity under hypoxia and dietary restriction [32]. FMOs contain the prosthetic group that binds FAD^+^ and utilize the cofactor NADPH together with O_2_ to catalyze substrate oxygenation, generating NADP^+^ as a by-product [33]. Animals subjected to INT1-specific *fmo-2* knockdown displayed a normal fasting-induced INS-7 secretion, but exhibited reduced basal INS-7 secretion and impaired refeeding response (Fig 6L). Given that NADPH/NADP^+^ ratio is a central determinant of cellular redox state and energy metabolism [34], this result further supports the notion that regulation of INS-7 secretion in INT1 cells is coupled to energy-sensing metabolic pathways.

Together, these results demonstrate that INT1 cells integrate immune and metabolic cues. By regulating pathogen avoidance and survival through lectin genes and modulating INS-7 secretion via energy-responsive genes such as *aakg-4* and *fmo-2,* INT1 cells function as a specialized intestinal subset that senses energy states and coordinates gut peptide secretion to maintain organismal homeostasis.

### Pyruvate is crucial for regulating INS-7 peptide secretion from INT1 cells

We were intrigued by the findings with *aakg-4* and *fmo-2*, which suggested that cellular energy status may govern the refeeding response of INS-7 secretion. Notably, the effect does not appear to be mediated by changes in intestinal fat content, as lipid levels remain unaltered during acute fasting and refeeding (Fig 1B and 1C). These observations led us to hypothesize that INT1 cells may integrate specific sensory cues to reset INS-7 secretion in response to food availability. To explore this possibility, we first evaluated the contribution of mechanosensory inputs. Fasted worms refed with 0.5 μm latex beads, comparable in size to *E. coli* [35, 36], failed to restore INS-7 secretion to basal levels, whereas refeeding with OP50 fully normalized INS-7 secretion (Fig 7A), suggesting that mechano-stimulus alone was not sufficient to induce the refeeding response of INS-7 secretion. Because *C. elegans* can detect secreted molecules from bacteria and trigger a stress response [37], we next exposed fasted animals to cell-free OP50 culture supernatant [37, 38]. Supernatant alone likewise failed to reset INS-7 secretion (Fig 7A). A combination of beads and supernatant was similarly ineffective (Fig 7A). In contrast, UV-killed OP50 completely reinstated basal INS-7 secretion (Fig 7A), indicating that chemicals or nutrients from bacteria, rather than mechano-stimulus or secreted molecules, mediate the refeeding response.

**Fig. 7.**
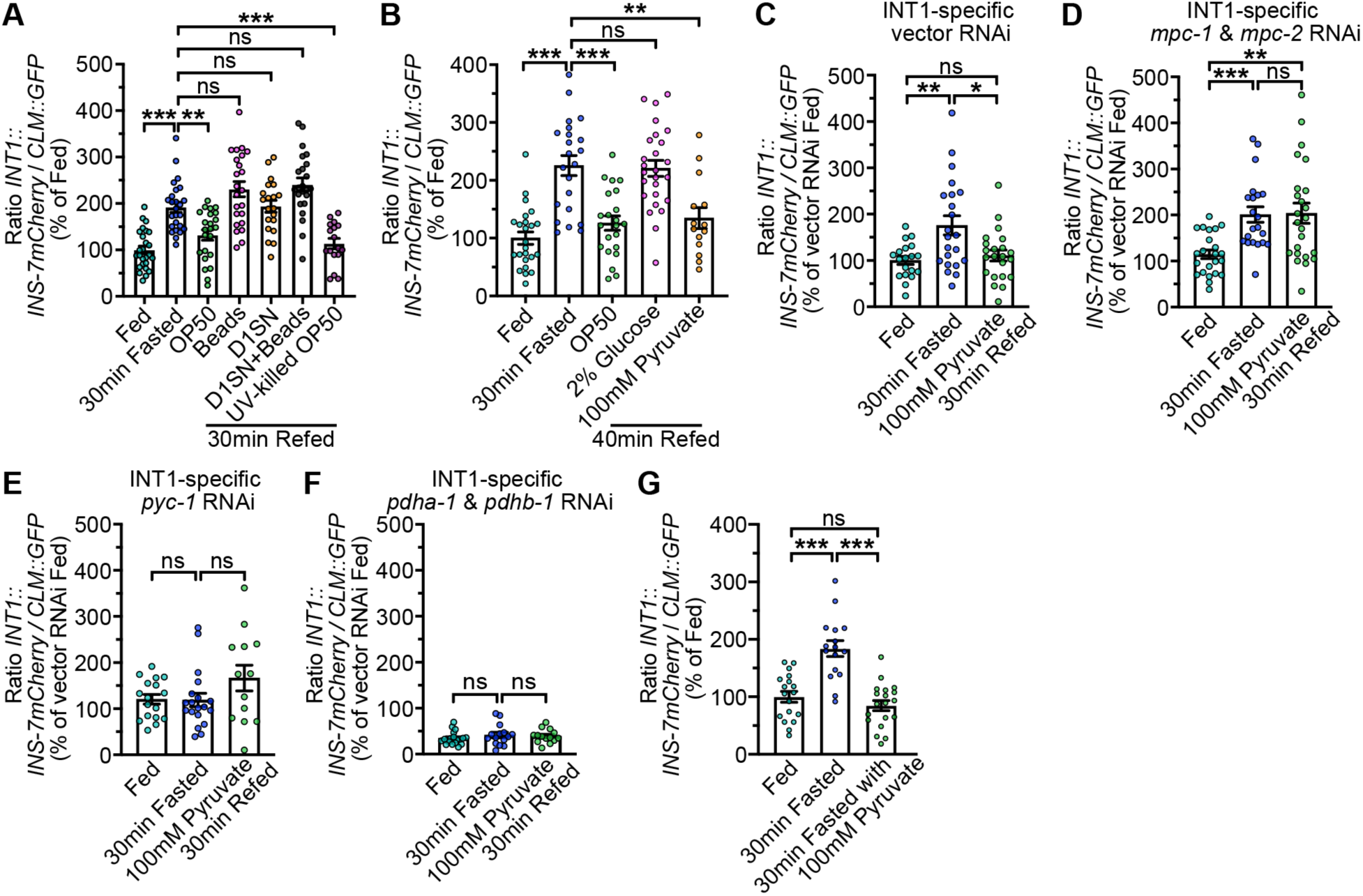
Pyruvate is critical for maintaining INS-7 secretion dynamics in INT1 cells. (A) INS-7 secretion dynamics during fasting and re-feeding with OP50, latex beads, Day 1 OP50 supernatant (D1SN), Day 1 OP50 supernatant with latex beads (D1SN+Beads), and UV-killed OP50 were determined at the indicated time points. The intensity of INS-7mCherry fluorescence within a single coelomocyte was quantified and normalized to the area of CLM::GFP expression for each time point. Data are expressed as a percentage of the normalized INS-7mCherry fluorescence intensity of wild-type fed animals ± SEM (n=20-26). **p<0.01, ***p<0.001, ^ns^p>0.05 by one-way ANOVA (Dunnett T3). (B) INS-7 secretion dynamics during fasting and re-feeding with OP50, 2% glucose and 100 mM pyruvate were determined at the indicated time points. The intensity of INS-7mCherry fluorescence within a single coelomocyte was quantified and normalized to the area of CLM::GFP expression for each time point. Data are expressed as a percentage of the normalized INS-7mCherry fluorescence intensity of wild-type fed animals ± SEM (n=14-26). **p<0.01, ***p<0.001, ^ns^p>0.05 by one-way ANOVA (Dunnett T3). (C-F) INS-7 secretion dynamics in INT1-specific vector RNAi, *mpc-1* and *mpc-2* RNAi, *pyc-1* RNAi, *pdha-1* and *pdhb-1* RNAi-treated animals during fasting and re-feeding with 100 mM pyruvate were determined at the indicated time points. The intensity of INS-7mCherry fluorescence within a single coelomocyte was quantified and normalized to the area of CLM::GFP expression for each time point. Data are expressed as a percentage of the normalized INS-7mCherry fluorescence intensity of vector RNAi fed animals ± SEM (n=13-24). *p<0.05, **p<0.01, ***p<0.001, ^ns^p>0.05 by one-way ANOVA (Dunnett T3). (G) INS-7 secretion dynamics during fasting and fasting with the presence of 100 mM pyruvate were determined at the indicated time points. The intensity of INS-7mCherry fluorescence within a single coelomocyte was quantified and normalized to the area of CLM::GFP expression for each time point. Data are expressed as a percentage of the normalized INS-7mCherry fluorescence intensity of fed animals ± SEM (n=16-19). ***p<0.001, ^ns^p>0.05 by one-way ANOVA (Dunnett T3).

Next, we set out to pinpoint the bacterial nutrient or chemical most critical for regulating INS-7 secretion during refeeding. Based on the requirement of *aakg-4* and *fmo-2* for INS-7 secretion dynamics, we tested whether restoring cellular energy states after fasting could return INS-7 secretion to baseline. Glucose serves as a primary cellular energy source, undergoing glycolysis to generate pyruvate, which in turn fuels downstream metabolic pathways to produce energy. However, refeeding fasted worms with 2% glucose, a concentration considered a high-glucose diet for *C. elegans* [47], failed to normalize INS-7 secretion, which remained elevated (Fig 7B). In contrast, remarkably, refeeding with 100 mM pyruvate, a dose effective in prior studies [47], fully recapitulated the refeeding response observed with OP50 (Fig 7B). This effect was dose-dependent (S9A Fig). Thus, pyruvate, a key molecule derived from glycolysis and a precursor metabolite for numerous biochemical processes, is sufficient to substitute for bacterial food in restoring INS-7 secretion to baseline after fasting.

After entering the cell, pyruvate can be converted to lactate or alanine in the cytosol or transported into the mitochondrial matrix via the mitochondrial pyruvate carrier (MPC) [39, 40] to enter the citric acid cycle, which generates energy [41]. INT1-specific RNAi of the two *C. elegans* MPC genes (*mpc-1* and *mpc-2*) abolished the pyruvate-induced restoration of INS-7 secretion (Fig 7C and 7D), demonstrating that mitochondrial import of pyruvate is required. Within the matrix, pyruvate is converted to acetyl-CoA by pyruvate dehydrogenase (*pdha-1* and *pdhb-1* in *C. elegans*) or to oxaloacetate by pyruvate carboxylase (*pyc-1* in *C. elegans*). INT1-specific knockdown of either pathway suppressed fasting-induced INS-7 secretion and abolished secretion dynamics (Fig 7E and 7F). Since chronic *pdha-1* and *pdhb-1* knockdown lowered basal INS-7 secretion level, we acutely inhibited pyruvate dehydrogenase during refeeding with the specific inhibitor 3-fluropyruvate (3-FP); this intervention prevented INS-7 secretion from returning to basal levels (S9B Fig), suggesting that the conversion of pyruvate to acetyl-CoA is required for the refeeding response of INS-7 secretion.

If intracellular pyruvate is the crucial determinant of INS-7 secretion, we predicted that maintaining pyruvate levels in the INT1 cells during fasting should suppress the fasting-induced INS-7 secretion. Indeed, worms fasted on NGM agar supplemented with 100 mM pyruvate showed no induction of INS-7 secretion (Fig 7G), suggesting pyruvate alone is sufficient to suppress fasting-induced INS-7 secretion. Collectively, these findings identify mitochondrial pyruvate metabolism as the key signal sensed by INT1 enteroendocrine cells to couple nutritional availability with the fluctuation in INS-7 peptide secretion during fasting and refeeding.

## Discussion

In the human intestine, both epithelial and immune cell compositions shift dramatically along the anterior-posterior axis, with region-specific cellular neighborhoods for epithelial, stromal, and immune cells contributing to distinct physiological functions [42]. In contrast, the *C. elegans* intestine comprises only 20 epithelial cells and lacks resident immune cells, raising the question of whether spatially defined molecular and functional heterogeneity also exists in this simplified but powerful model system. Our findings, together with prior observations, support the view that *C. elegans* intestinal subsets are molecularly and functionally distinct. The data show that the most pronounced difference between INT1 and the rest of the intestine at the RNAseq level, is the expression of stress response genes in INT1. Although there are some nuanced differences, the prevalence of stress response genes persists across feeding and fasting conditions. This difference, combined with the evidence that INT1 cells secrete the enteroendocrine peptide INS-7 [6] (Fig 6) is strongly reminiscent of the mammalian enteroendocrine cells, which also secrete peptides and show strong expression of stress response genes [43, 44]. We suggest that this category term reflects not only a canonical stress response, but also a broader response to shifts in the luminal environment, which INT1 cells are anatomically poised to detect well before the absorption of nutrients has begun further down the intestine (INT2-9). Thus, we believe INT1 cells are a newly defined enteroendocrine cell-type within the *C. elegans* intestine.

In the present study, we utilized INT1- and INT2-9-specific promoters to perform TRAP-Seq analyses under fasting and refeeding conditions. While comparisons between INT1 and INT2-9 cells revealed marked differences, this approach likely underrepresents heterogeneity among the INT2-9 cells themselves. Prior studies demonstrated that posterior intestinal cells initiate the rhythmic defecation motor program [45, 46], and our dataset identified *npr-28*, which is specifically expressed in the posterior intestinal cells INT7-9 (Fig 2J), further underscoring molecular diversity across intestinal segments. Future identification of promoters with more restricted expression within the INT2-9 subset will be crucial for functionally resolving the spatial map of the *C. elegans* intestine.

We previously identified INT1 as enteroendocrine cells that secrete the gut peptide INS-7 preferentially during acute food withdrawal to suppress neuronally stimulated fat loss [6]. Additional lines of evidence also point to the specialization of the anterior-most INT1 cells. First, acid phosphatase staining for intestinal brush border could not be detected in INT1 cells [15, 47]. Second, Methyl red staining showed strong red staining of acidic vacuoles in INT1 cells. From INT2 to INT9 cells, vacuoles were stained gradually from orange-yellow (neutral-alkaline) to yellow (alkaline) with methyl red [47], suggesting the pH of intracellular vacuoles is lower in INT1 cells. Third, ultrastructural analysis highlighted enrichment of rough ER, Golgi, and mitochondria in INT1, consistent with a secretory function, whereas INT2-9 cells preferentially accumulate yolk and lipid vacuoles for storage [48]. Fourth, components of a cullin-RING ubiquitin ligase complex critical for thermotolerance and the intracellular pathogen response (IPR), including CUL-6 and SKR-5, were found to be specifically expressed in INT1 cells [49], suggesting that INT1 cells may play an integral role in proteostasis responses to intracellular pathogen infection. Collectively, these findings establish INT1 as a specialized cell-type that integrates endocrine, stress, and immune functions within the *C. elegans* intestine.

We uncovered pyruvate as a crucial metabolite that functions as a true surrogate for bacterial food to regulate the secretion of the endogenous insulin antagonist INS-7 [6]. Pyruvate must enter the TCA cycle to elicit this effect, suggesting that either actual metabolism of pyruvate is required, or that downstream metabolites may also act as mediators of this process. In other systems, TCA cycle intermediates such as acetyl-CoA, itaconate, and L-2-hydroxyglutarate (L-2-HG) modulate immune responses [50], while succinate and fumarate regulate hypoxia sensing upon cytosolic release [51]. A similar mechanism may underpin the communication between mitochondria and the nucleus in *C. elegans*, exemplified by the translocation of ATFS-1 during the activation of the mitochondrial unfolded protein response [52]. Thus, pyruvate metabolism may provide a metabolic checkpoint that couples nutrient availability to enteroendocrine signaling.

Many bacterial species including *Y. enterocolitica*, *C. freundii*, *E. aerogenes* and *E. coli* excrete pyruvate into the extracellular environment during nutrient-replete conditions [53, 54] and reabsorb extracellular pyruvate during nutrient insufficiency [55, 56]. Thus, the presence of extracellular pyruvate may serve as an indicator of the nutrient abundance in bacterial environments, a sensing mechanism that *C. elegans* may have co-evolved by leveraging the INT1 cells to couple environmental conditions with host physiology. It was reported that *C. elegans* exhibits chemotaxis to volatile pyruvic acid via AWC neurons [57]. However, the molecular mechanisms underlying both INT1 and neuronal sensing of pyruvate remain uncharacterized. Thus both intracellular and extracellular bacterial pyruvate appear to regulate the metabolism or behavior of *C. elegans*, indicating that pyruvate is a crucial signaling molecule.

In summary, we expect our TRAP-Seq datasets presented here to provide the scientific community with valuable resources for investigating the dynamic translatomes of specific subsets of intestinal cells in response to fasting and refeeding. More broadly, the discovery of cell-type heterogeneity within the *C. elegans* intestine highlights the evolutionary conservation of spatially organized intestinal biology and lays a foundation for future mechanistic studies of metabolic and immune regulation in the intestine.

### Limitations of the TRAP-Seq in the current study

Not all previous studies have demonstrated a strong correlation between mRNA expression levels and protein abundance [58–60]. TRAP-Seq provides a more precise assessment of gene expression by detecting actively translating mRNAs. Specifically, TRAP-Seq better reflects the translational responses under various conditions, such as temperature fluctuations or fasting. Moreover, by employing tagged ribosomal proteins, TRAP-Seq enables the selective purification of polysomes without the need for cell dissociation or sorting, thereby reducing RNA contamination from neighboring cells or tissues.

To obtain sufficient ribosome-mRNA complexes for RNA sequencing, we generated strains with an integrated extrachromosomal array expressing HA-tagged RPL22 in either INT1 or INT2-9 cells. While this strategy enables robust affinity purification, the elevated expression of HA-tagged RPL22 in this system could potentially introduce unanticipated effects on transcript utilization or translational bias. Nevertheless, because all analyses rely on internal referencing, the differential gene expression signatures remain valid. In addition, the random integration of multi-copy transgenes into unknown genomic loci may alter the expression of adjacent genes, leading to additional variability. Importantly, we did not observe any changes in growth rate, fertility or fecundity in the TRAP-seq transgenic lines, suggesting that potential confounding effects are minimal.

TRAP has traditionally tagged proteins of the 60S large ribosomal subunit, including RPL10A [10, 11, 61–64] and RPL22 [13, 65–67], to enable tissue- or cell type-specific capture of polysome-mRNA complexes followed by RNA sequencing. A key distinction among previous studies lies in the choice of ribosomal protein for tagging. Both RPL10A and RPL22 are evolutionarily conserved ribosomal proteins. In *C. elegans*, RPL10A [64, 68, 69] and RPL22 [13, 67, 70] have been successfully used for tagging in TRAP-Seq experiments, yielding robust results. However, recent findings in mammalian systems indicate that RPL10A is significantly sub-stoichiometric in polysomes of mouse embryonic stem cells [71], suggesting it is not associated with all actively translating mRNAs. Similarly, in *C. elegans*, RPL10A shows limited expression in the germline and weak association with ribosomes, impairing the co-purification of translating mRNAs from the germline [70]. These observations suggest that tagging RPL10A in TRAP experiments may not capture the complete translatome in specific cell types or tissues. Further, *Nousch et al.* recently demonstrated that RPL-4 and RPL-9 are more effective than RPL10A and RPL22 for purifying polysome-mRNA complexes from germ cells in *C. elegans* [70], suggesting that the choice of ribosomal proteins for tagging in TRAP-Seq experiments may vary depending on the tissue of interest. Specifically, certain ribosomal proteins may offer improved specificity and efficiency for isolating tissue-specific mRNAs. Importantly, the RPL22-based TRAP in our experiments achieved robust immunoprecipitation efficiency (about 40% of input) for intestinal polysomal mRNAs (S4 table), thereby validating our approach. Future studies may further investigate whether alternative ribosomal proteins could provide enhanced isolation of translating mRNAs in different tissues including the intestine.

## Materials and methods

### *C. elegans* maintenance and strains

Worms were cultured on nematode growth medium (NGM) agar plates with *Escherichia coli* strain OP50 at 20°C as described [72]. Bristol strain N2 was obtained from the *Caenorhabditis* Genetic Center (CGC) and used as the wild-type strain. All transgenic strains used in this study are listed in S1 Table. Worms were synchronized for Oil Red O staining by hypochlorite treatment, after which hatched L1 larvae were seeded on plates with the appropriate bacteria; worms were synchronized for INS-7 peptide secretion assay, PA14 avoidance and slow killing assays by letting gravid adult worms lay eggs on plates with the appropriate bacteria for 1.5 hr. All experiments were performed on day 1 adults.

### Acute and chronic fasting

For the fasting experiments, synchronized day 1 adult worms were removed from the OP50 seeded NGM plates with M9 buffer, washed 3 times to remove remaining OP50 then placed on the unseeded NGM plates for 30 minutes (acute fast) or 180 minutes (chronic fast). For the refeeding experiments, fasted worms were removed from the fasting plate with M9 buffer then placed onto OP50 seeded plates for 30 minutes (acute fast/refeeding) or 90 minutes (chronic fast/refeeding).

### Oil Red O staining

Oil Red O staining was performed as described [73]. Worms were harvested with phosphate-buffered saline (PBS) and incubated on ice for 10 minutes. Worms were fixed as described [73]. Following fixation, worms were stained in filtered Oil Red O (Thermo Scientific) working solution (60% Oil Red O in isopropanol: 70% water) overnight. Approximately 2000 worms were fixed and stained for all conditions within one experiment. We visually examined about 100 worms on slides and then blindly chose 20 worms for imaging. All experiments were repeated at least three times.

### Image acquisition and quantitation

Oil Red O-stained worms were imaged using 10X objective on a Zeiss Axio Imager microscope. Images were acquired with the Software AxioVision (Zeiss). Lipid droplet staining in the first four pairs of intestinal cells was quantified using ImageJ software (NIH). All reported results were consistent across biological replicates. Fluorescent images of mNeonGreen reporters were acquired with software AxioVision (Zeiss) using a 10X objective on a Zeiss Axio Imager microscope. Fluorescent images of reporters for INS-7 secretion were acquired with software AxioVision (Zeiss) using a 20X objective on a Zeiss Axio Imager microscope. The images were taken at the indicated time point or immediately after treatment, such as refeeding. The first pair of coelomocytes was imaged. mCherry fluorescence intensity in one of the two imaged coelomocytes was quantified and normalized to the surface area of the coelomocyte (*Punc-122::GFP*) as previously described and validated [74]. Within each experiment, at least 15 worms from each condition were quantified using ImageJ software (NIH).

### Cloning and transgenic strain construction

For generating *Pges-1ΔB::rpl-22-3xHA* and *Ppho-1::rpl-22-3xHA* constructs, promoters and *rpl-22-3xHA* were amplified with standard PCR techniques and subcloned into donor vectors using Gateway Cloning Technology^TM^ (Life Technologies). All plasmids for NeonGreen reporters were generated by Gibson Assembly^TM^. Primers used for cloning in this study are listed in S3 Table. Transgenic strains were generated by injecting plasmids into the germline of wild-type worms followed by visual selection of co-injection markers (*Pmyo-2::mCherry*, *Pges-1ΔB::GFP* or *Ppho-1::mCherry*) under the fluorescence microscope. The INS-7 secretion line was previously developed and validated [6]. For *INT1::TRAP* strain, N2 worms were injected with 10 ng/μL of the *Pges-1ΔB::rpl-22-3xHA* plasmid, 20 ng/μL of the *Pges-1ΔB::GFP* plasmid, and 70 ng/μL of an empty vector to bring the final concentration of injection mix to 100 ng/μL. For *INT2-9::TRAP* strain, N2 worms were injected with 25 ng/μL of the *Ppho-1::rpl-22-3xHA* plasmid, 50 ng/μL of the *Ppho-1::mCherry* plasmid, and 25 ng/μL of an empty vector to bring the final concentration of injection mix to 100 ng/μL. A transgenic line with high transmission rate and consistent expression was integrated using the Stratalinker UV Crosslinker 2400 (Stratagene) and backcrossed four times before experimentation. For other microinjections, we injected the animals with 30 ng/μL of the desired plasmid, 10 ng/μL of the *Pmyo-2::mCherry* and 60 ng/μL empty vector to maintain a final injection mix concentration of 100 ng/μL. Two lines were selected for experimentation based on the transmission rate and consistency of expression.

### RNAi

RNAi experiments were performed as described [75, 76]. Carbenicillin-IPTG plates were seeded with HT115 bacteria containing the empty or the relevant RNAi clone four days before seeding larvae.

### PA14 avoidance assay

The PA14 avoidance assay was performed as previously described [77]. *Pseudomonas aeruginosa* strain PA14 was cultured by inoculating a single colony into 2 mL of LB broth and incubating at 37 °C for 6 hr. 20 μL of the culture was then seeded onto 3.5 cm slow-killing (SK) plates and incubated at 37 °C for 15 hr. For each assay, 30 synchronized Day 1 gravid adults (containing 6-10 eggs) grown on RNAi bacteria were cleaned and transferred to the agar surface outside the PA14 lawn. The number of animals on and off the PA14 lawns was scored after 8 h. Three plates were used per RNAi treatment in each experiment.

### PA14 slow killing assay

PA14 slow killing assay was performed as described [30]. Briefly, overnight cultures of *Pseudomonas aeruginosa* strain PA14 were seeded onto slow-killing (SK) plates and incubated at 37°C for 24 hr, followed by 25°C for an additional 24 hr. Synchronized animals at the closed vulva stage subjected to RNAi treatment were transferred to 3.5 cm SK plates seeded with PA14 (30 animals per plate), with three plates per RNAi treatment. Plates were maintained at 25°C, and worm survival was monitored until all animals had died. Survival curves were analyzed using GraphPad Prism, and statistical significance was determined by the log-rank test.

### Translating ribosome affinity purification

The protocol was adapted from reference [13] and described in detail in S1 Protocol. Briefly, transgenic worms containing the Ribo-TRAP constructs were made to drive expression of *rpl-22-3xHA* in INT1 and INT2-9. Age-matched transgenic worms for each experimental condition were washed, flash-frozen, and lysed to release ribosomes and cellular components in the presence of cycloheximide to pause translation as well as RNAase and protease inhibitors. Immunoprecipitation of ribosomes from our desired tissue was performed with a monoclonal anti-HA antibody and protein G-beads. Nascent RNA was eluted from ribosomes with beta mercaptoethanol (BMP). Percent IP RNA compared to input, on average less then 1%, was used to determine if the IP was successful.

### RNA purification

RNA from the IP, flowthrough, and input was purified with Agilent Absolutely RNA Nanoprep Kit^TM^ according to the manufacturer’s instructions including the on-column DNase treatment. Final RNA isolates were eluted in 15 μl and used to make diluted aliquots for sequencing, validation, and QC.

### RNA quality assessment

We tested for RNA degradation and quality by running the input, flowthrough, and IP samples on an Agilent Bioanalyzer chip (nano or pico, depending on concentration). Because IP samples had a small concentration it was difficult to accurately measure the RNA integrity number (RIN) in all conditions, but the quality of RNA for these unknown samples could be extrapolated from the flowthrough and input samples. In all conditions the average RIN across sample types was greater than 7.5 indicating a RNA integrity sufficient for sequencing, and in the majority of conditions it was greater than 9 indicative of high-quality samples.

### RNAseq library prep, sequencing, and alignment

Total RNA samples were prepared into RNAseq libraries using the NEBNext® Ultra™ Directional RNA Library Prep Kit for Illumina® following the manufacturer’s recommended protocol. Briefly, for each sample 100ng total RNA was polyA-selected, converted to double stranded cDNA followed by fragmentation and ligation of sequencing adapters. The libraries were then PCR amplified 12 cycles using barcoded PCR primers, purified and size selected using AMPure XP Beads before loading onto an Illumina NextSeq500 for 75 base single read sequencing. Reads were aligned to *C. elegans* genome (WBcel235/r67) using STAR (version 2.6.1d) [78], and gene counts were produced using featureCounts (version 1.6.4) [79]. On average, 10.6 million unique single-end (SE) reads aligned per INT1 sample and 19.5 million unique SE reads per INT2-9 sample, differences were due to the larger RNA input for INT2-9 samples. Data quality was assessed using FastQC (version 0.11.8) [80]. Raw RNA-seq FASTQ files and a gene count matrix are available on GEO (GSE306302). Bioinformatic analysis was performed using R programming language (version 4.4.2) (R Development Core Team, 2015) and RStudio integrated development environment (version 2024.12.1) (Team R, 2018). Plots were generated using ggplot2 R package (version 3.5.2).

### Differential expression analysis

Differential expression analysis was performed using the DESeq2 R package (version 1.46.0) [81] which models the gene expression counts with a negative binomial distribution while adjusting for library size and dispersion. Differentially expressed genes were identified as having adjusted p-value less then 0.05 using a Benjamini-Hochberg false discovery rate correction. DESeq2 variance stabilized counts were used to generate normalized count matrices that were used for plotting expression data of individual genes and those averaged across WGCNA modules. PCA was performed using the top 500 variable genes from variance stabilized counts output in DESeq2. Statistical significance of PCA correlation was determined with a Pearson’s correlation for 2 factor comparisons (line) or an ANOVA for multi-factor comparisons (metabolic stress). Plot ellipses comparing groups were drawn with ggplot2 stat_ellipse() function which computes the mean and covariance of the PC1 and PC2 for each line to build the ellipse at a 95% confidence level.

### WGCNA

WGCNA was performed with the WGCNA R package (version 1.73) [16] to identify modules of co-expressed genes and their associations with external traits in the analysis. Prior to analysis, samples with low expression values were removed to reduce variation associated with batch, a 30 min fasting and 30 min refeeding sample were removed from the *Pges-1ΔB* experiment based on this criteria. DESeq2 variance stabilized counts were used as the input and only genes in the 95^th^ percentile of variance were kept to reduce noise (n = 2,338). A soft-thresholding power of 10 was selected using the scale-free topology criterion to approximate scale-free network topology. A signed adjacency matrix was constructed and transformed into a topological overlap matrix (TOM), followed by hierarchical clustering to detect modules of highly correlated genes. Modules were identified using dynamic tree cutting with a minimum module size of 30 genes and subsequently merged based on eigengene similarity (cut height = 0.25). Module eigengenes (MEs) were correlated with external traits (e.g., line and fasting condition), and Pearson correlation coefficients along with associated p-values were computed. Modules showing significant trait associations were further investigated for biological relevance and functional enrichment. Module GO enrichment analysis was performed using the ClusterProfiler R package version (version 4.14.6)[82]. For GO analysis, we reported terms with FDR < 0.1. The test set of module genes was compared against the background set of genes in the WGNCA analysis passing the variance criteria.

### Statistics

Wild-type animals were included as controls for every experiment. Error bars represent standard error of the mean (SEM). Student’s *t*-test and one-way ANOVA were used as indicated in the Fig legends. All statistical analyses were performed in GraphPad Prism 9.3 (GraphPad Software). Appropriate multiple comparison corrections were used for ANOVAs.

## Acknowledgements

This work was supported by research grants to S.S. from the NIH/NIDDK (R01 DK124706) and NIH/NIA (R01 AG056648). This work was supported by the resources and staff at the Scripps Research Genomics Core Facility. C.L. was supported by a Dorris Scholar Award from the Dorris Neuroscience Center, The Scripps Research Institute. Elements of Fig 1F were created with BioRender.com.

## FIGURES and FIGURE LEGENDS

**S1 Fig.**
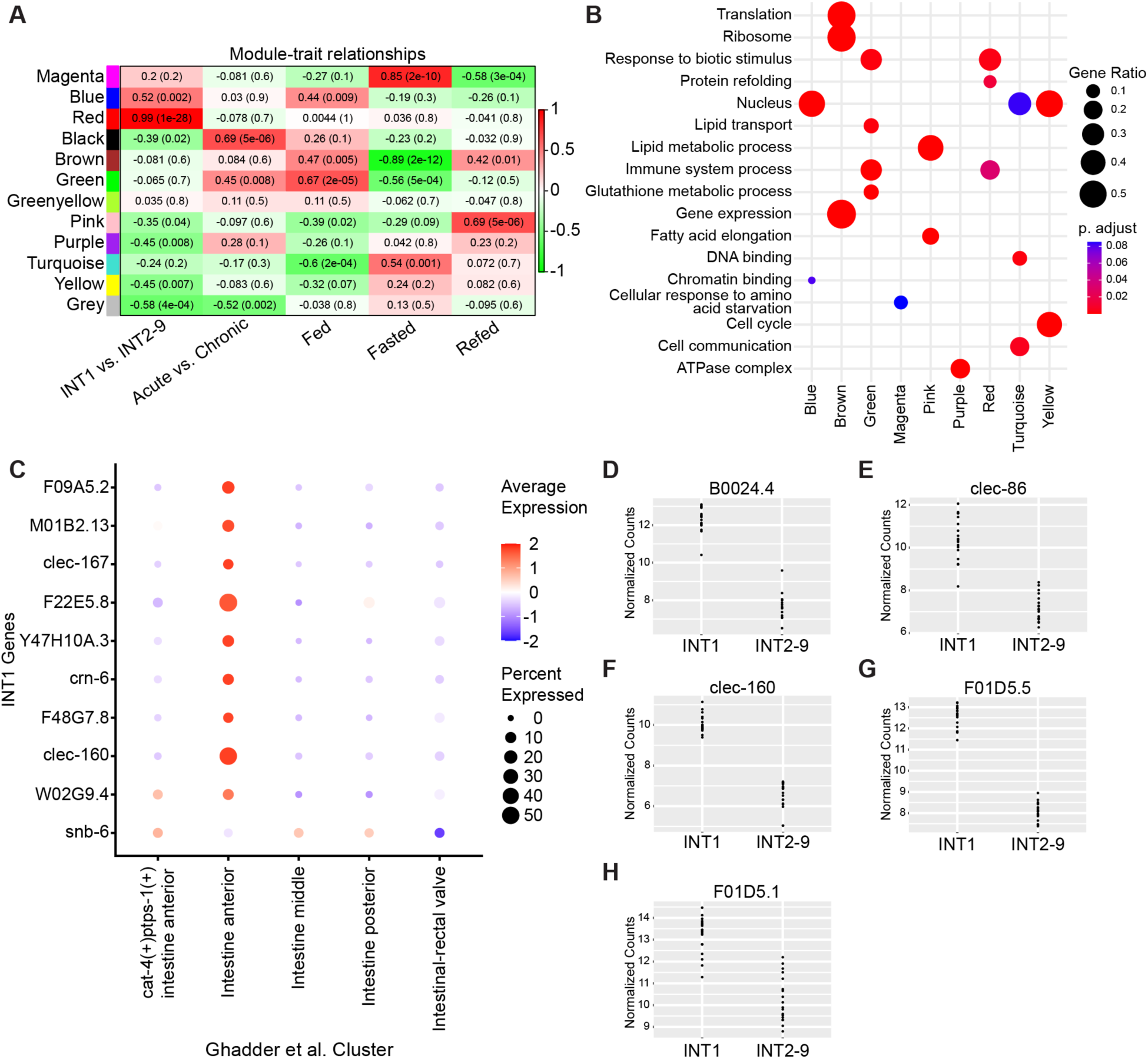
The genes differentially expressed in INT1 cells. (A) Module-trait relationships for selected conditions in WGCNA. Rows indicate the WGCNA module and the columns comparative traits. The color indicates the strength of the Pearson’s correlation and bracketed number the significance. The color red represents a positive correlation; the color green represents a negative correlation. (B) GO enrichment of selected modules in the WGCNA analysis. Size indicates the strength of the significance and color the significance. (C) Dotplot showing increased expression in the anterior intestine clusters from Ghadder et al. for genes upregulated in the INT1 vs INT2-9 Fed datasets. We selected the top 10 significant (FDR < 0.05) genes ranked by effect size that were expressed in at least 5% of anterior intestinal cells to show. Size indicates the percent of cells expressing the gene, and color indicates the scaled effect size. (D-H) Expression profiles of *B0024.4, clec-86, clec-160, F01D5.5* and *F01D5.1* under all conditions in INT1 and INT2-9 cells.

**S2 Fig.**
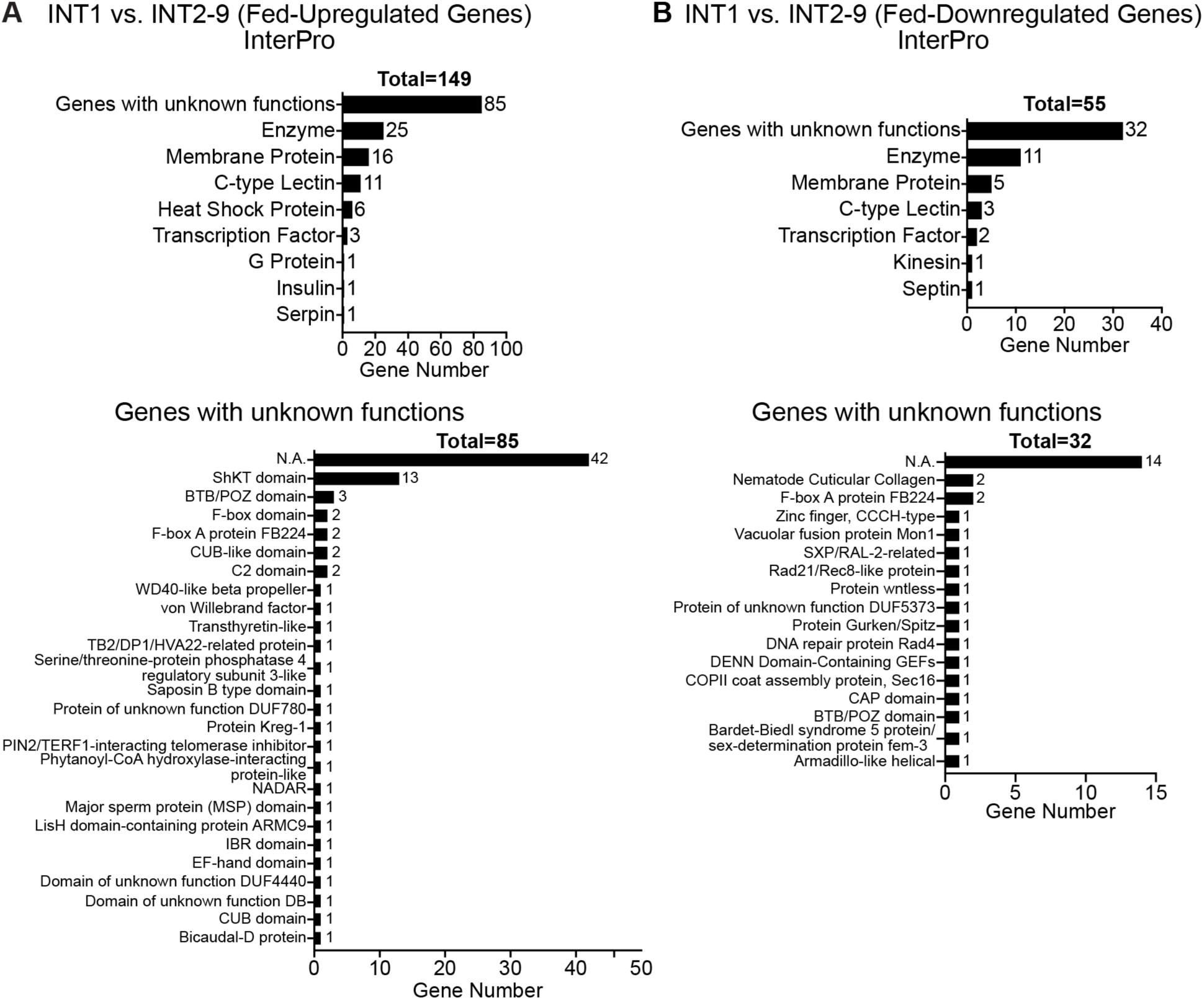
InterPro annotations of differentially expressed genes for INT1 Fed versus INT2-9 Fed. (A, B) Bar charts of the upregulated or downregulated genes categorized by the protein domains predicted by InterPro for INT1 fed versus INT2-9 fed.

**S3 Fig.**
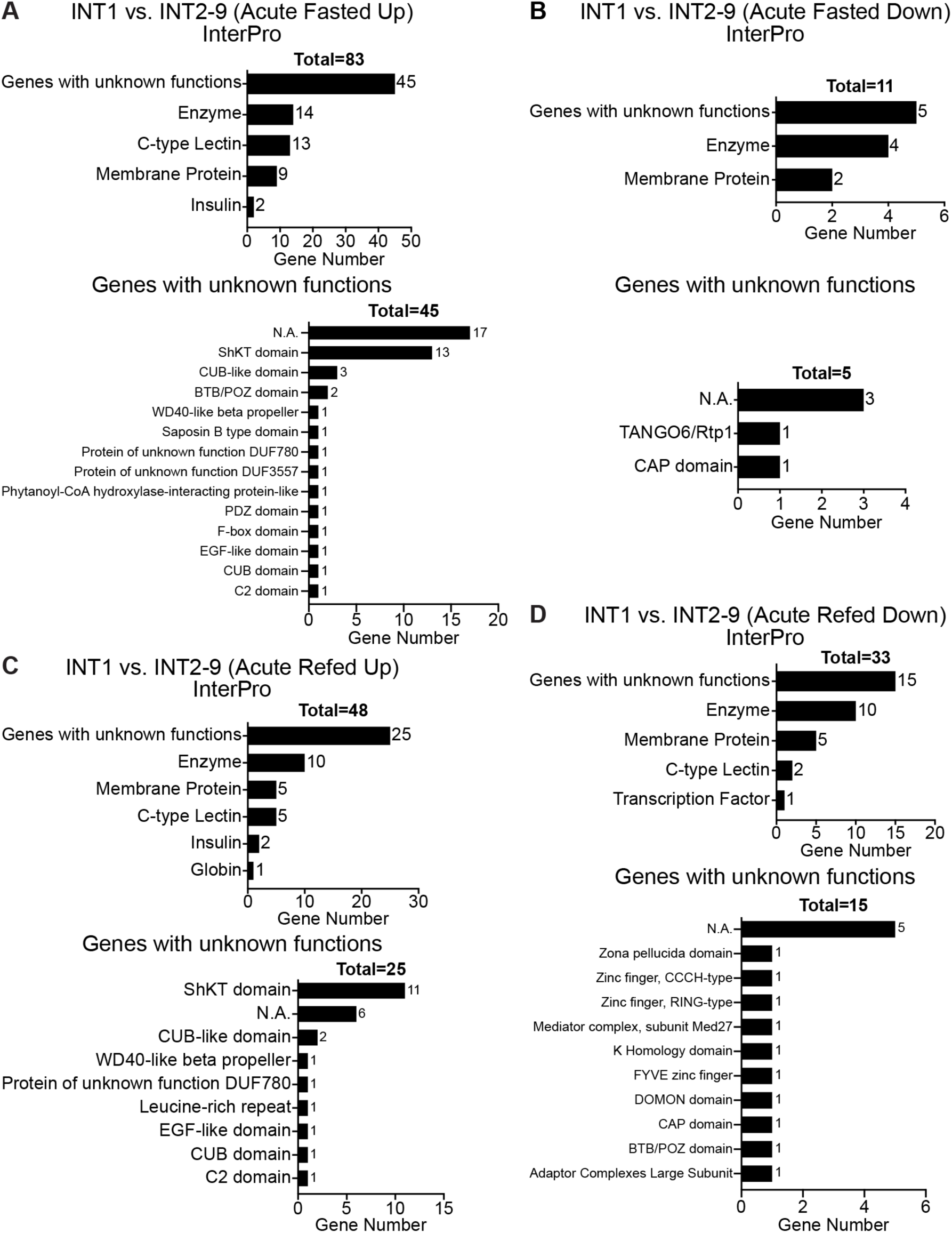
InterPro annotations of differentially expressed genes for INT1 versus INT2-9 under acute conditions. (A, B) Bar charts of the upregulated or downregulated genes categorized by the protein domains predicted by InterPro for INT1 30 min fasted versus INT2-9 30 min fasted. (C, D) Bar charts of the upregulated or downregulated genes categorized by the protein domains predicted by InterPro for INT1 30 min refed versus INT2-9 30 min refed.

**S4 Fig.**
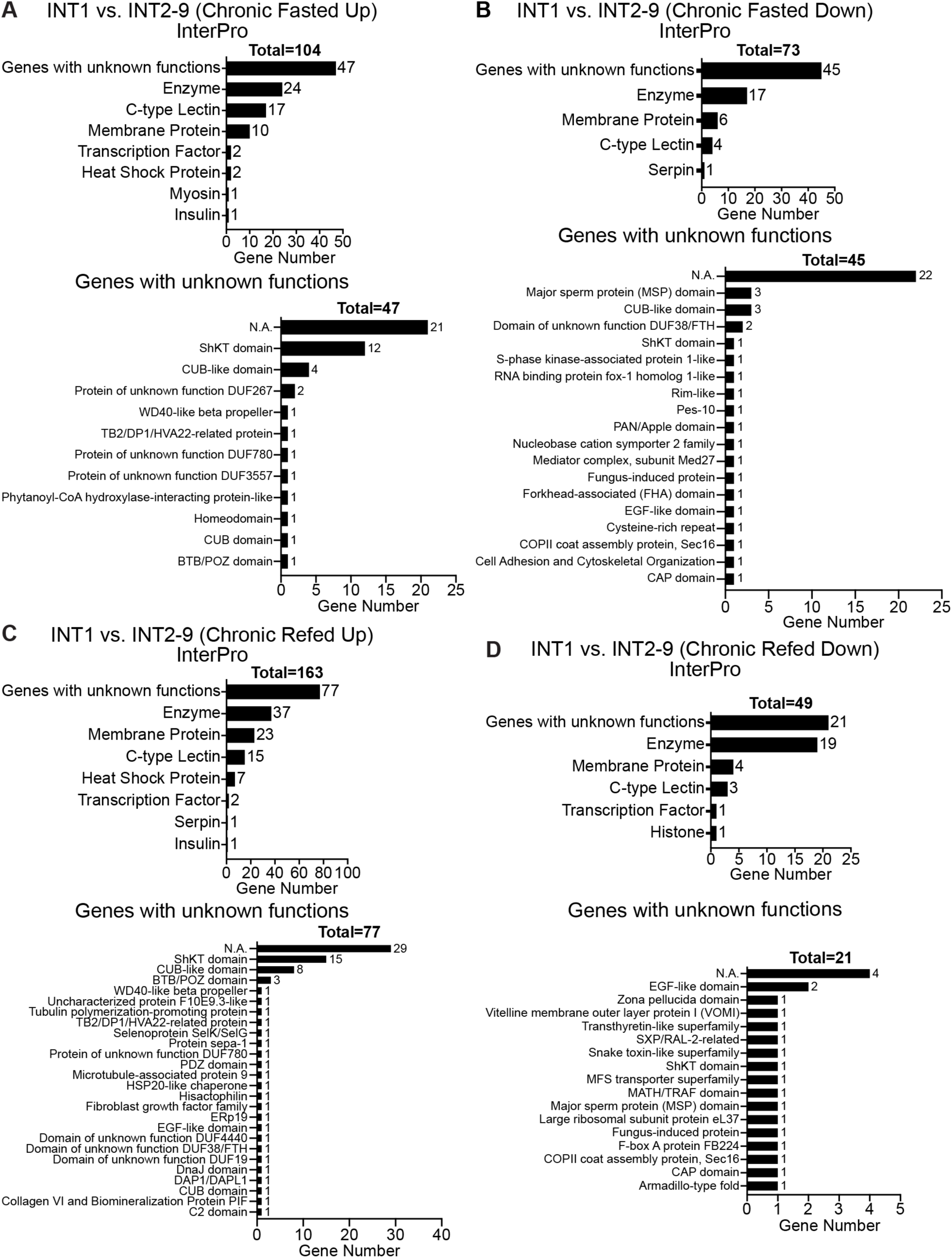
InterPro annotations of differentially expressed genes for INT1 versus INT2-9 under chronic conditions. (A, B) Bar charts of the upregulated or downregulated genes categorized by the protein domains predicted by InterPro for INT1 180 min fasted versus INT2-9 180 min fasted. (C, D) Bar charts of the upregulated or downregulated genes categorized by the protein domains predicted by InterPro for INT1 90 min refed versus INT2-9 90 min refed.

**S5 Fig.**
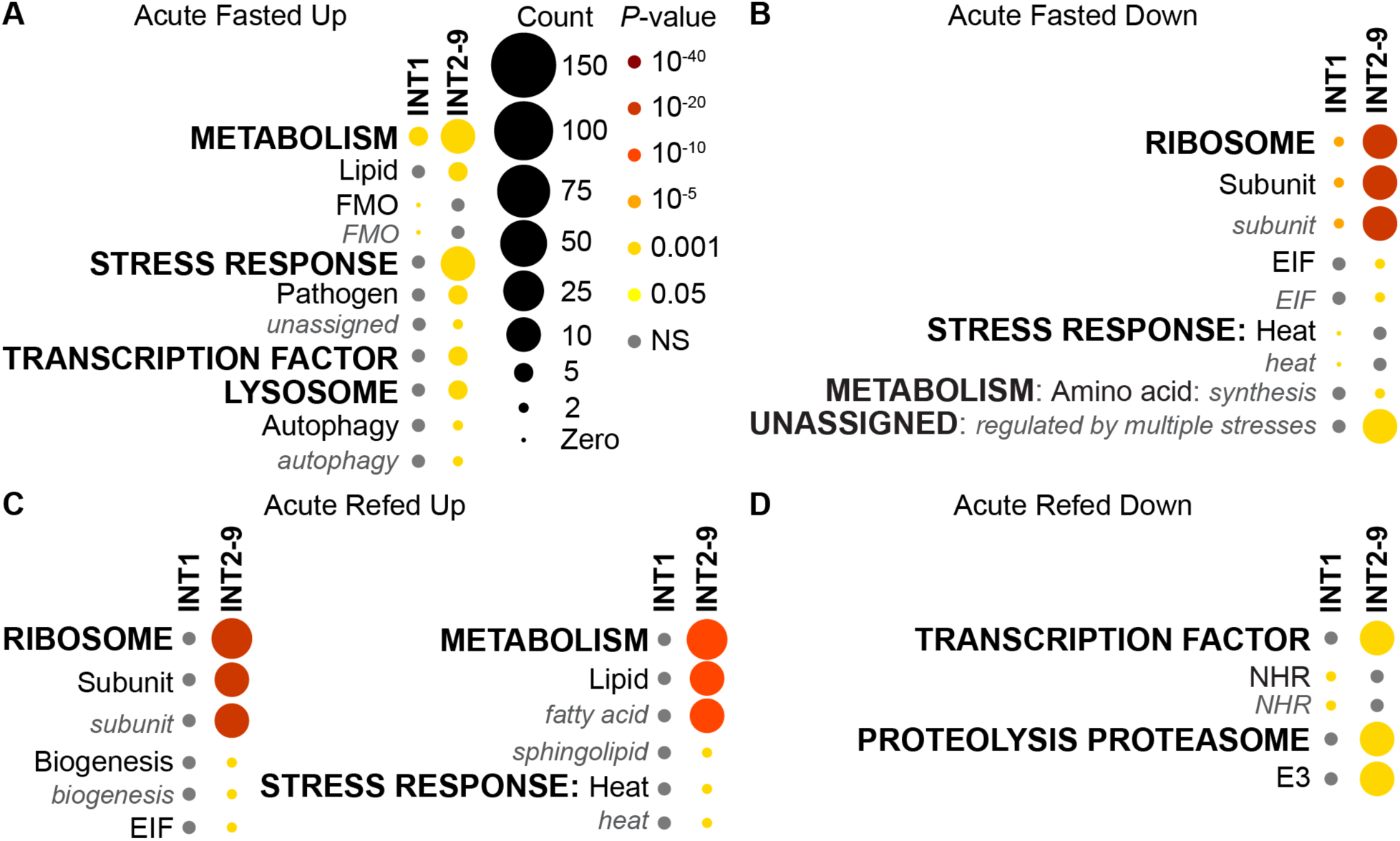
WormCat annotations of differentially expressed genes under acute fasting and refeeding conditions for INT1 and INT2-9 cells. (A, B) WormCat visualization of categories enriched in differentially expressed genes in INT1 cells and INT2-9 cells under acute fasting condition. (C, D) WormCat visualization of categories enriched in differentially expressed genes in INT1 cells and INT2-9 cells under acute refeeding condition. Categories 1 are all bold uppercase; Categories 2 are capitalized; Categories 3 are gray italics. The size of the bubbles indicates the gene counts in the category and the color of the bubble represents the adjusted p-value.

**S6 Fig.**
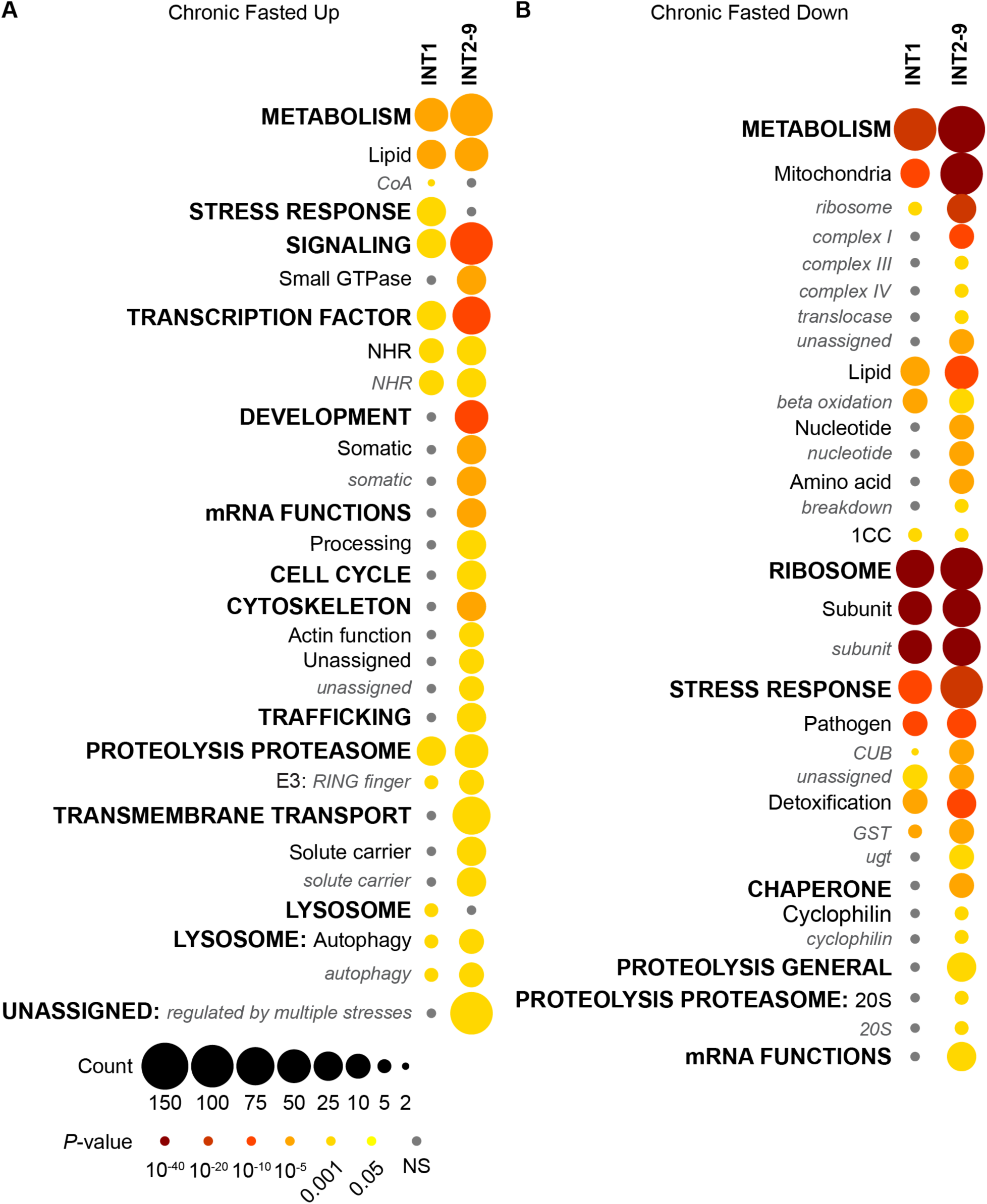
WormCat annotations of differentially expressed genes under chronic fasting condition for INT1 and INT2-9 cells. (A, B) WormCat visualization of categories enriched in differentially expressed genes in INT1 cells and INT2-9 cells under chronic fasting condition. Categories 1 are all bold uppercase; Categories 2 are capitalized; Categories 3 are gray italics. The size of the bubbles indicates the gene counts in the category and the color of the bubble represents the adjusted p-value.

**S7 Fig.**
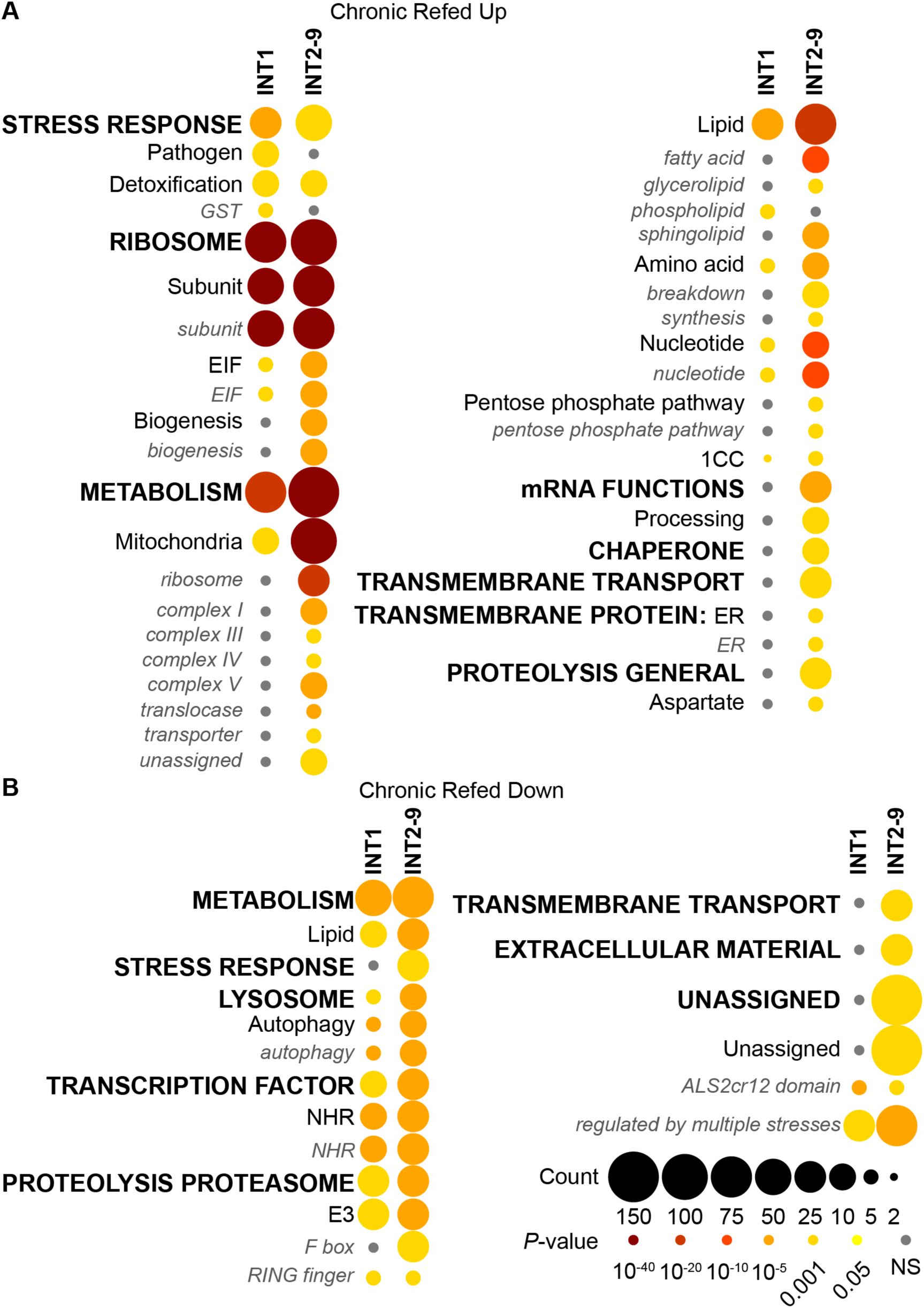
WormCat annotations of differentially expressed genes under chronic refeeding condition for INT1 and INT2-9 cells. (A, B) WormCat visualization of categories enriched in differentially expressed genes in INT1 cells and INT2-9 cells under chronic refeeding condition. Categories 1 are all bold uppercase; Categories 2 are capitalized; Categories 3 are gray italics. The size of the bubbles indicates the gene counts in the category and the color of the bubble represents the adjusted p-value.

**S8 Fig.**
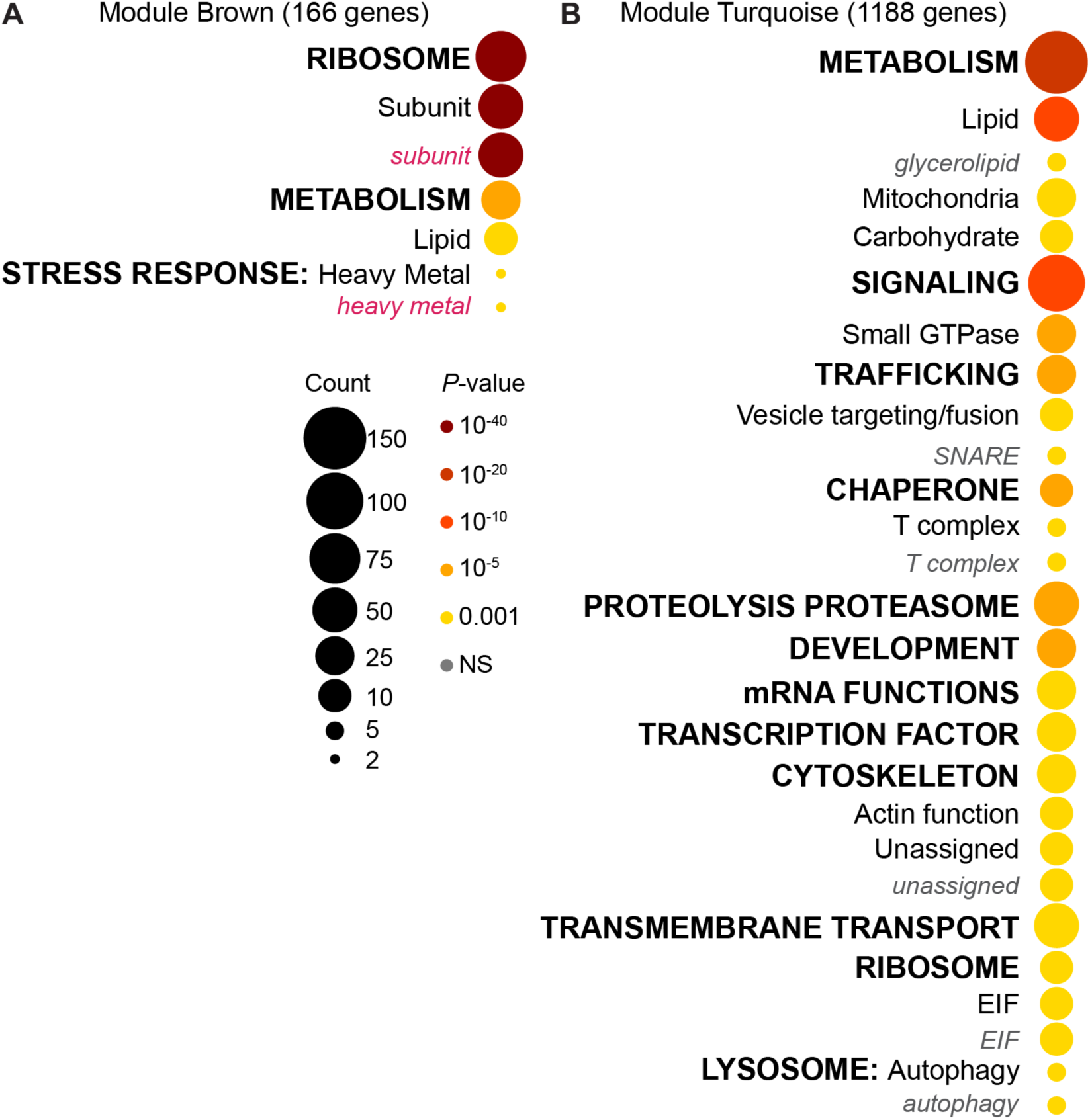
WormCat annotations of undulating genes in INT1 and INT2-9 cells. (A, B) WormCat visualization of categories enriched in genes of module brown and turquoise. Categories 1 are all bold uppercase; Categories 2 are capitalized; Categories 3 are gray italics. The size of the bubbles indicates the gene counts in the category and the color of the bubble represents the adjusted p-value.

**S9 Fig.**
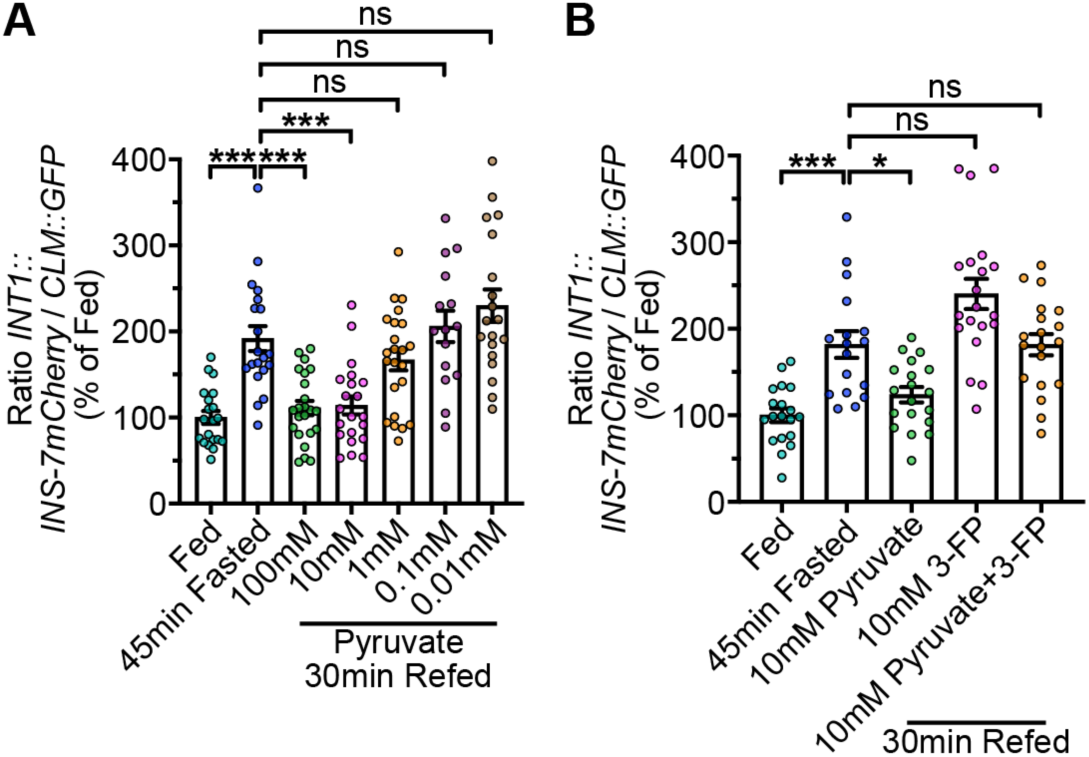
The role of pyruvate in refeeding responses of INS-7 secretion from INT1 cells. (A) INS-7 secretion dynamics during fasting and re-feeding with different concentrations of pyruvate (0.01 mM - 100 mM) were determined at the indicated time points. The intensity of INS-7mCherry fluorescence within a single coelomocyte was quantified and normalized to the area of CLM::GFP expression for each time point. Data are expressed as a percentage of the normalized INS-7mCherry fluorescence intensity of wild-type fed animals ± SEM (n=15-23). ***p<0.001, ^ns^p>0.05 by one-way ANOVA (Dunnett T3). (B) INS-7 secretion dynamics during fasting and re-feeding with 10 mM pyruvate, 10 mM 3-FP, and 10 mM pyruvate and 3-FP were determined at the indicated time points. The intensity of INS-7mCherry fluorescence within a single coelomocyte was quantified and normalized to the area of CLM::GFP expression for each time point. Data are expressed as a percentage of the normalized INS-7mCherry fluorescence intensity of wild-type fed animals ± SEM (n=17-20). *p<0.05, ***p<0.001, ^ns^p>0.05 by one-way ANOVA (Dunnett T3).

---

**Supplementary Table 1:** *C. elegans* strains used in this study.

| Strains used in this study | Source | Identifier | Additional Information |
| --- | --- | --- | --- |
| *N2*; *ssrls1248*[*Pges-1ΔB::rpl-22-3xHA*]; *ssrls1191*[*Pges-1ΔB::GFP*] | This paper | SSR1578 | Integrated. Backcrossed 4X. |
| *N2*; *ssrls1191*[*Pges-1ΔB::GFP*]; *ssrls1210*[*Ppho-1::mCherry*] | This paper | SSR1582 | Integrated. Backcrossed 4X. |
| *N2*; *ssrls1251*[*Ppho-1::rpl-22-3xHA*]; *ssrls1210*[*Ppho-1::mCherry*] | This paper | SSR1605 | Integrated. Backcrossed 4X. |
| *sid-1(qt9)*; *ssrls1220*[*Pges-1ΔB::sid-1::GFP*]; *ssrls1240*[*Pges-1ΔB::ins-7mCherry*]; *ssrls615*[*Punc-122::GFP*] | PMID: 37961386 | SSR1634 |  |
| *N2*; *ssrEx1317*[*Pclec-160::mNeonGreen*]; *ssrEx1031*[*Pmyo-2::mCherry*] | This paper | SSR1701 |  |
| *N2*; *ssrEx1319*[*Pclec-86::mNeonGreen*]; *ssrEx1031*[*Pmyo-2::mCherry*] | This paper | SSR1702 |  |
| *N2*; *ssrEx1318*[*PB0024.4::mNeonGreen*]; *ssrEx1031*[*Pmyo-2::mCherry*] | This paper | SSR1703 |  |
| *N2*; *ssrEx1323*[*PF01D5.5::mNeonGreen*]; *ssrEx1031*[*Pmyo-2::mCherry*] | This paper | SSR1705 |  |
| *N2*; *ssrEx1324*[*Pnpr-28::mNeonGreen*]; *ssrEx1031*[*Pmyo-2::mCherry*] | This paper | SSR1706 |  |
| *N2*; *ssrEx1325*[*PF01D5.1::mNeonGreen*]; *ssrEx1031*[*Pmyo-2::mCherry*] | This paper | SSR1713 |  |

**Supplementary Table 2:** Plasmids used in this study.

| Plasmids used in this study | Source | Identifier |
| --- | --- | --- |
| *Pges-1ΔB::GFP* | This paper | pSS1191 |
| *Ppho-1::mCherry* | This paper | pSS1210 |
| *Pges-1ΔB::rpl-22-3xHA* | This paper | pSS1248 |
| *Ppho-1::rpl-22-3xHA* | This paper | pSS1251 |
| *Pclec-160::mNG::clec-160 3'UTR* | This paper | pSS1317 |
| *PB0024.4::mNG::B0024.4 3'UTR* | This paper | pSS1318 |
| *Pclec-86::mNG::clec-86 3'UTR* | This paper | pSS1319 |
| *PF01D5.5::mNG::F01D5.5 3'UTR* | This paper | pSS1323 |
| *Pnpr-28::mNG::npr-28 3'UTR* | This paper | pSS1324 |
| *PF01D5.1::mNG::F01D5.1 3'UTR* | This paper | pSS1325 |

**Supplementary Table 3:**
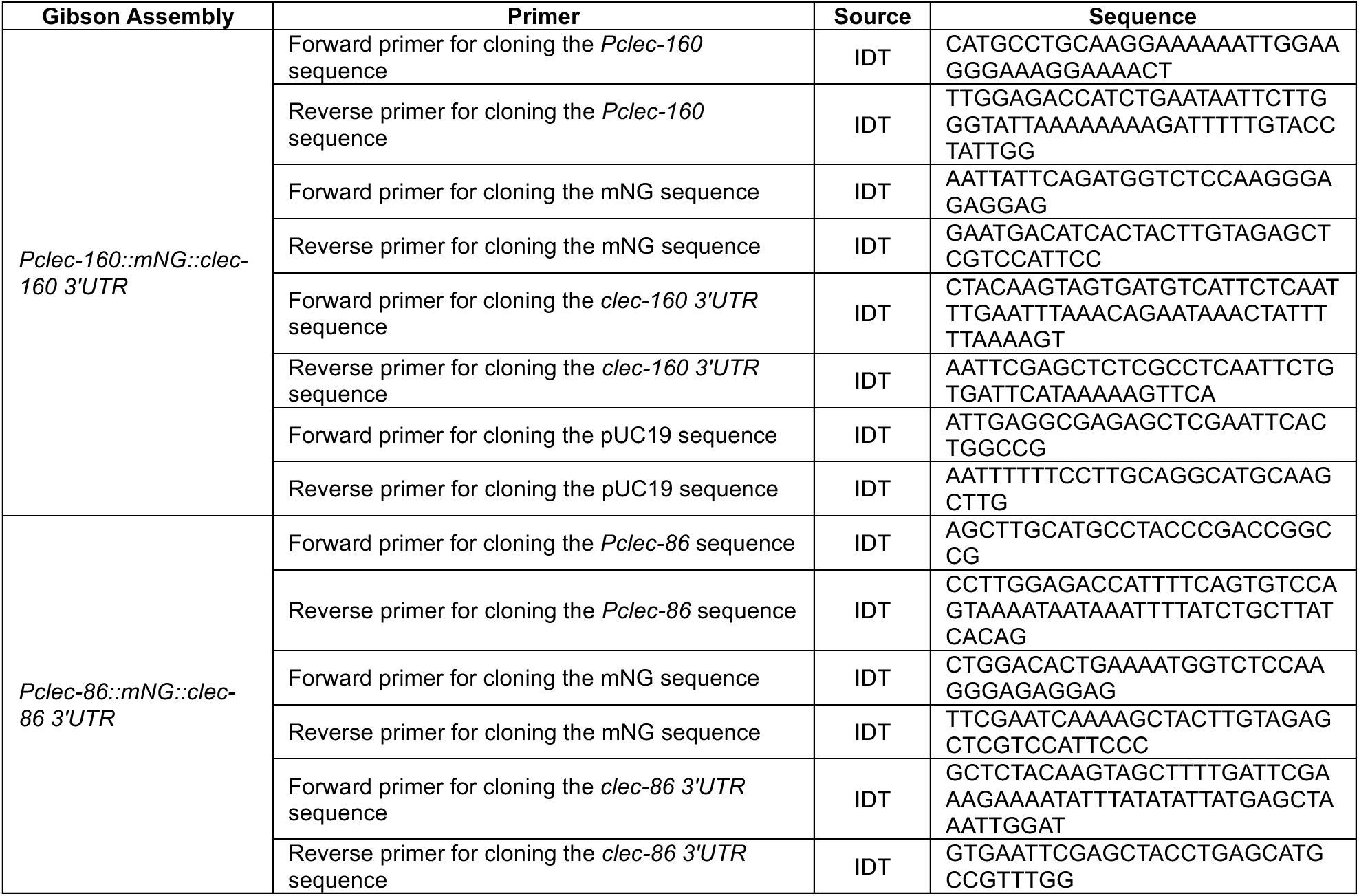
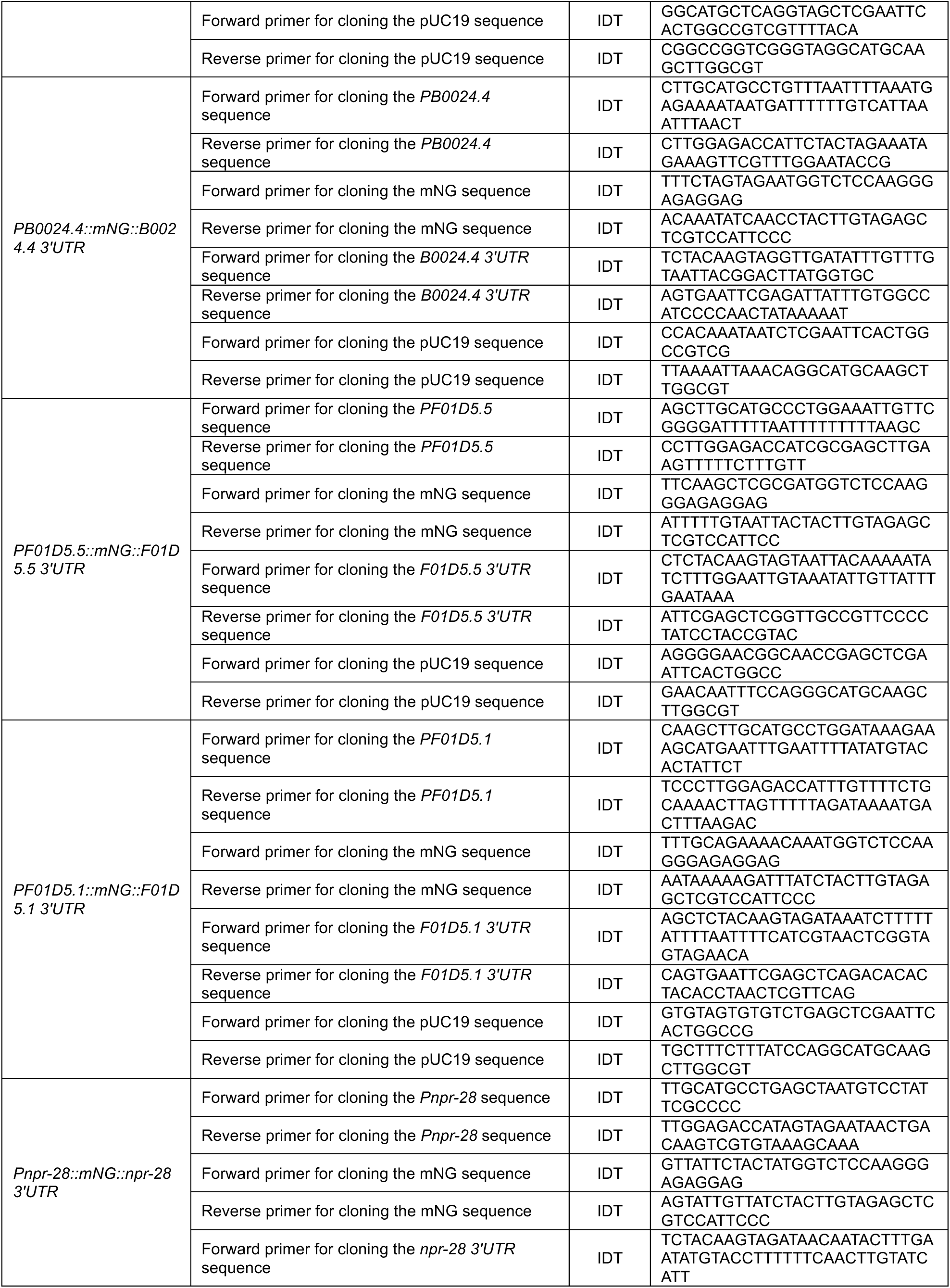
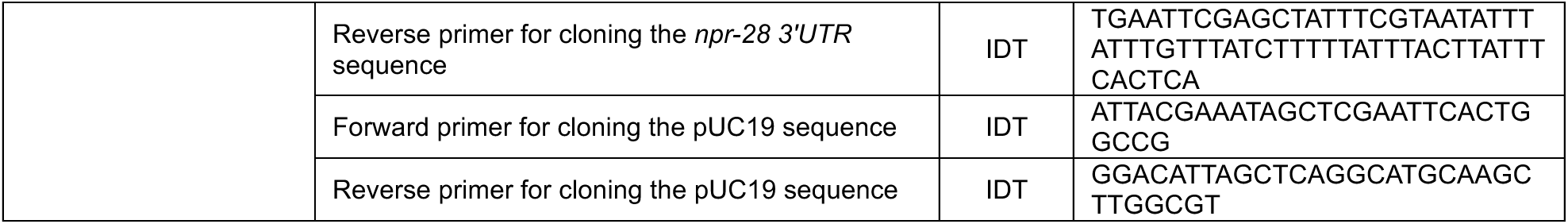
Cloning primers used in this study.

